# The relationship between PD-L1 and quiescence in melanocyte stem cell aging

**DOI:** 10.1101/2022.09.22.508528

**Authors:** Joseph W. Palmer, Kyrene M. Villavicencio, Misgana Idris, Dominique Weddle, Fabian V. Filipp, NISC Comparative Sequencing Program, William J. Pavan, Melissa L. Harris

**Affiliations:** Department of Biology, University of Alabama at Birmingham, Birmingham, AL, USA; Institute of Computational Biology, Helmholtz Zentrum München, Ingolstädter Landstr. 1, 85764 Neuherber; NIH Intramural Sequencing Center, National Human Genome Research Institute, National Institutes of Health, Bethesda, MD, USA; Genetic Disease Research Branch, National Human Genome Research Institute, National Institutes of Health, Bethesda, MD, USA

**Keywords:** aging, quiescence, PD-L1, melanocyte stem cell, hair graying

## Abstract

A central aspect of life-long stem cell function in slow cycling stem cells is the proper regulation of cellular quiescence. How the quiescent state is achieved, whether all quiescent cells are equivalent, and if the quiescent stem cell pool changes with age are all questions that remain unanswered. Using quiescent melanocyte stem cells (qMcSC) as a model, we found that stem cell quiescence is neither a singular nor static process and can be heterogeneous. As one example of this heterogeneity, we show that a portion of qMcSCs expresses the immune checkpoint protein PD-L1 at the cell membrane (PD-L1^mem+^), PD-L1^mem+^ qMcSCs are better retained with age, and that the aged quiescent McSC pool is transcriptomically more deeply quiescent. Collectively these findings demonstrate that PD-L1 expression is a physiological attribute of quiescence in McSCs and PD-L1^mem+^ quiescent stem cells may be good targets for reactivation in the context of aging.

## Introduction

Healthy aging depends on the proper and persistent function of tissue-specific stem cell populations throughout the body (Ahmed et al., 2017; Oh et al., 2014). Numerous studies have shown that a variety of genetic and environmental factors can contribute to dysfunction in stem cell maintenance and self-renewal resulting in premature tissue aging (Cheung et al., 2015; Harris et al., 2018; Inomata et al., 2009; Uno and Nishida, 2016). The dormant state of cellular quiescence (G_0_) is the primary method by which premature proliferative exhaustion of stem cell populations is prevented with age (Tümpel and Rudolph, 2019). Although often described simply as a state of reversible non-proliferation, recent discoveries suggest that regulation of G_0_ is remarkably diverse and dynamic (Cho et al., 2019; Coller et al., 2006; Kwon et al., 2017; Urbán and Cheung, 2021). The G_0_ state allows stem cells to remain poised for reactivation. Upon receipt of specific activation cues, stem cells rapidly reenter the cell cycle to produce differentiated progeny to restore and maintain tissue homeostasis (Cheung and Rando, 2013; Velthoven and Rando, 2019). In fact, several genes are upregulated during G_0_ and knock-down studies have shown that these cell state-specific genes are required for preventing stem cell depletion suggesting that G_0_ is more tightly regulated than previously thought (Du et al., 2012; Mourikis et al., 2012; Zhou et al., 2018). Yet, few studies have focused on evaluating whether stem cells during G_0_ are molecularly equivalent and whether all G_0_ stem cells undergo age-related changes. We anticipate that the composition of the G_0_ stem cell pool changes with age and that these changes predict the regenerative capacity of that tissue in the context of aging.

Uncovering the extent and complexity of G_0_ within mammalian stem cell populations and determining whether age-associated changes in these molecular programs occur *in vivo* remains challenging. However, melanocyte stem cells (McSCs) residing within the hair follicle stem cell niche provide an ideal model to address this problem. McSCs are responsible for providing the differentiated, pigment-producing melanocytes that reside in the hair follicle bulge. These melanocytes deposit melanin into the growing hair follicle, which results in the coloration of our hair (Slominski et al., 1994, 2005). Gray hair is one of the most readily recognized, and outwardly visible, age-associated phenotypes and has been largely attributed to the gradual depletion of McSCs with age (Inomata et al., 2009; Nishimura et al., 2005). McSC activation and G_0_ are easily evaluated because these processes are inextricably tied to distinct stages of the hair cycle. The hair cycle consists of three main stages, growth (anagen), regression (catagen), and dormancy (telogen) and the timing of each of these stages is well-characterized in mice (Müller-Röver et al., 2001).

A highly enriched population of quiescent McSCs (qMcSCs) can be isolated from the telogen stage of the hair cycle using melanocyte-specific cell surface markers based on the fact that telogen hairs are devoid of differentiated melanocytes (Harris et al., 2018). This advantageous feature, inherent to the biology of McSCs, allowed us to uncover an underlying network of genes specifically upregulated during G_0_ and identify major biological processes associated with qMcSCs. Notably, we discovered that qMcSCs are enriched in genes associated with immune system processes including upregulation of the immune checkpoint protein, programmed death-ligand 1 (*Pd-l1*). Interestingly, expression of PD-L1 at the cell membrane (PD-L1^mem+^) marks only a subpopulation of qMcSCs *in vivo*, which reveals heterogeneity in the qMcSC pool. We also find that PD-L1^mem+^ is tied to the G_0_ state of non-tumorigenic melanocytic cells *in vitro* and that PD-L1^mem+^ expression increases as a function of G_0_ length. Lastly, we show that with age, the transcriptome of McSCs resembles that of a deeper G_0_ state and this is concurrent with changes in the aged McSC pool where PD-L1^mem-^ _neg_ qMcSCs are depleted while PD-L1^mem+^ qMcSCs are retained. The results of this study highlight the molecular changes that occur during G_0_ in young and aged G_0_ stem cells, links PD-L1^mem+^ expression to the G_0_ state, and points to increased G_0_ depth as a novel paradigm for stem cell aging.

## Results

### Defining an *in vivo* transcriptome regulating stemness and quiescence in McSCs

To evaluate the effects of aging on McSC G_0_ we first established an *in vivo* gene expression profile that distinguishes adult qMcSCs from their non-G_0_ McSC precursors or melanoblasts. Using tissue dissociation and flow cytometry, KIT+/CD45-melanocytic cells were isolated from the dermis of mice at two time points that are distinct developmentally and in their cell state (stem cell precursors versus stem cells, and actively proliferating versus G_0_) (**Figure 1a**). Cells were harvested at P0.5 to encompass proliferating melanoblasts colonizing the hair follicle and at 8 weeks to capture adult qMcSCs that reside in dormant telogen-stage hairs using methods similar to those described previously (Harris et al., 2018). These enriched cells were processed for RNA sequencing (RNA-seq) and differential gene expression analysis. Downstream analysis showed that 8705 genes were differentially expressed between these two cell populations (q-value < 0.05, absolute log2 fold-change (L2FC) > 0.5; **Supplemental File 1**). We interpret these differentially expressed genes (DEGs) to reflect the global shift in gene expression required for actively proliferating melanoblasts to successfully colonize the hair follicle stem cell niche and transition to the adult, G_0_ and stem cell state. As expected for qMcSCs, our RNA-seq data showed decreased expression in several cell cycle related genes (*Ki67, Cdk2*, and *Cdk4*) and melanogenesis-related genes (*Dct, Mitf, Pmel, Tyr*, and *Sox10*). Conversely, we found increased expression of known stemness genes including *Epas1, Hes1, Klf4, Klf10*, and *Sox9*, along with several genes associated with the regulation of G_0_ in other stem cell populations including, *Cdkn1a, Nfatc1, Col12a1*, and *Arid5a* (**Figure 1b**). To identify relevant molecular pathways that contribute to maintaining (i) the G_0_ state and/or (ii) the undifferentiated stemness properties of McSCs, we selected the outer 10% of DEGs upregulated in either melanoblasts or qMcSCs for further evaluation. This top 10% was based on the highest-ranked expression of the normalized mean of counts across all samples. This resulted in a short-list of what we refer to as the “core DEGs”. We used these core-DEGs to represent the proliferative (516, ‘Melanoblast Core’) and G_0_/stem (357, ‘Adult qMcSC Core’) transcriptomic signatures (**Figure 1c; Supplemental File 2**). The L2FC distribution showed that these highly expressed genes tend to have a lower overall L2FC and are often arbitrarily filtered out when using a standard 2-fold cut-off (**Figure 1c**). Yet, due to the highly expressed nature of these genes in relation to the total size of the transcriptome, this subset of core-DEGs has a high potential for biological significance. Further assessment of the core-DEGs respective to each cellular state showed well-known proliferation markers such as *Ki67* and *Top2a* to be upregulated by proliferating melanoblasts, and increased expression of protein homeostasis (*Hspa1a, Hspa1b*) and adhesion genes (*Cadm2, Cdh2, L1cam*,) in qMcSCs. To determine the biological processes associated with the transition of proliferating melanoblasts into qMcSCs, we evaluated the top 20 enriched biological processes using gene set enrichment analysis (GSEA). The results showed an overrepresentation of biological processes such as RNA processing and translation initiation to be associated with the proliferating melanoblasts, while regulation of cell proliferation, differentiation, and cell death were associated with qMcSCs (**Figure 1d; Supplemental File 2**). Altogether these results suggest that G_0_ in McSCs is not idle and is instead actively maintained by various genes involved in several biological processes that underlie this dormant state. Upregulation of G_0_-specific genes and pathways further demonstrate that G_0_ in qMcSCs is not simply reflective of the downregulation of all major cellular processes.

**Figure 1.**
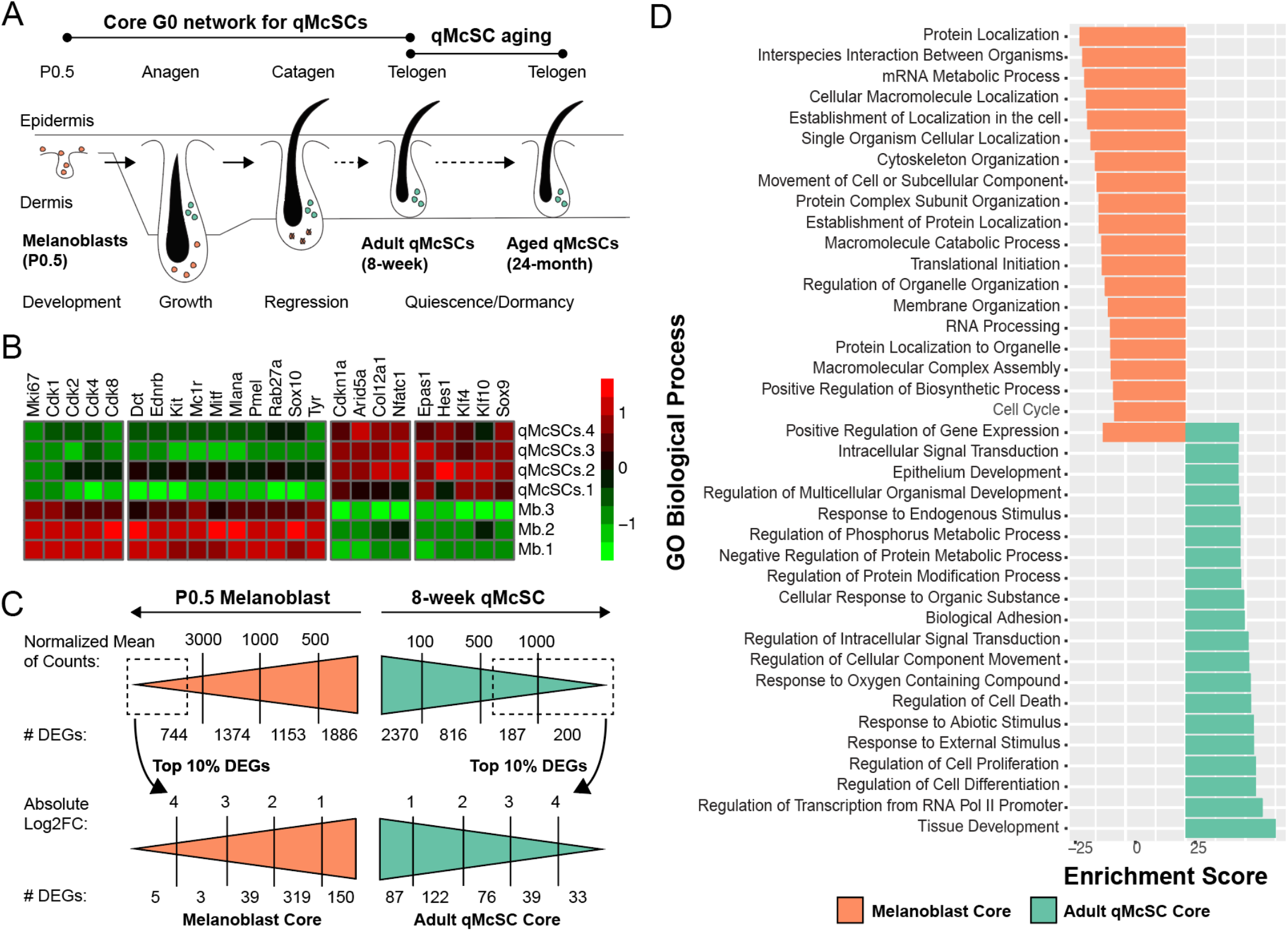
(A) Diagram depicting the three main stages of the hair cycle (anagen, catagen, and telogen) and the timepoints used for RNA-seq comparisons of melanocytic cells (P0.5 melanoblasts, 8-week adult qMcSCs, and 24-month aged qMcSCs). (B) Heatmap showing differential expression of the cell cycle, lineage, stemness, and G0-associated genes between P0.5 melanoblasts and 8-week adult qMcSCs. (C) The distribution of mean normalized counts of statistically significant DEGs (q-value < 0.05, L2FC > 0.5) identified between P0.5 melanoblasts and 8-week adult qMcSCs. The top 10% of highly expressed DEGs were included as part of the melanoblast and adult qMcSC core gene sets. (D) Clustered bar graph of the top 20 enriched biological processes associated with the two sets of core DEGs.

### Elevated *Pd-l1* expression is associated with qMcSCs

To identify novel mediators of G_0_ in qMcSCs we first identified hub genes with elevated expression based on cellular state. Similar to the role of transcription factors, hub genes are defined as having an above-average number of interaction partners within the genome and simultaneously influence the expression patterns of these interacting genes (Fox et al., 2011; Liu et al., 2019). Thus, hub genes with elevated expression are likely to play a role in regulating major biological processes regulating cell state. Identification of hub genes within the core DEGs for the two time points was performed using the Enrichr database. Core DEGs were overlapped with previously defined hub genes within the mammalian genome, with hub genes defined as those genes found to have greater than 120 protein-protein interactions (PPIs) (Chen et al., 2013a; Kuleshov et al., 2016). We found a total of 39 and 9 of these hub genes to be associated with the proliferative (P0.5 melanoblasts) and G_0_ (8-week qMcSCs) cellular states, respectively (**Figure 2a**). Additionally, we identified 36 transcription factors (TFs) in qMcSCs, 4 of which are also considered hub genes, that we predict act as upstream regulators to drive or maintain G_0_ and stemness within this cell population (**Supplemental File 2**). Previous studies have shown that altered expression in 8/36 (23.5%) of these TFs are linked to pigmentary defects in multiple organisms including mice and humans with *Atf3, Bcl6, Epas1, Hes1, Myc, Sox9, Trp63*, and *Vdr* showing increased expression in qMcSCs (Baxter et al., 2018). Further experiments testing the necessity of each hub gene or TFs will elucidate the biological roles these genes have on regulating stemness or G_0_ in McSCs. Of particular interest was a cluster of TFs known to be key regulators of immune system processes including the hub gene *Stat3* as well as, *Stat2, Irf1*, and *Irf9*. Until now, the relevance and role of immune genes in qMcSCs have not been evaluated.

**Figure 2.**
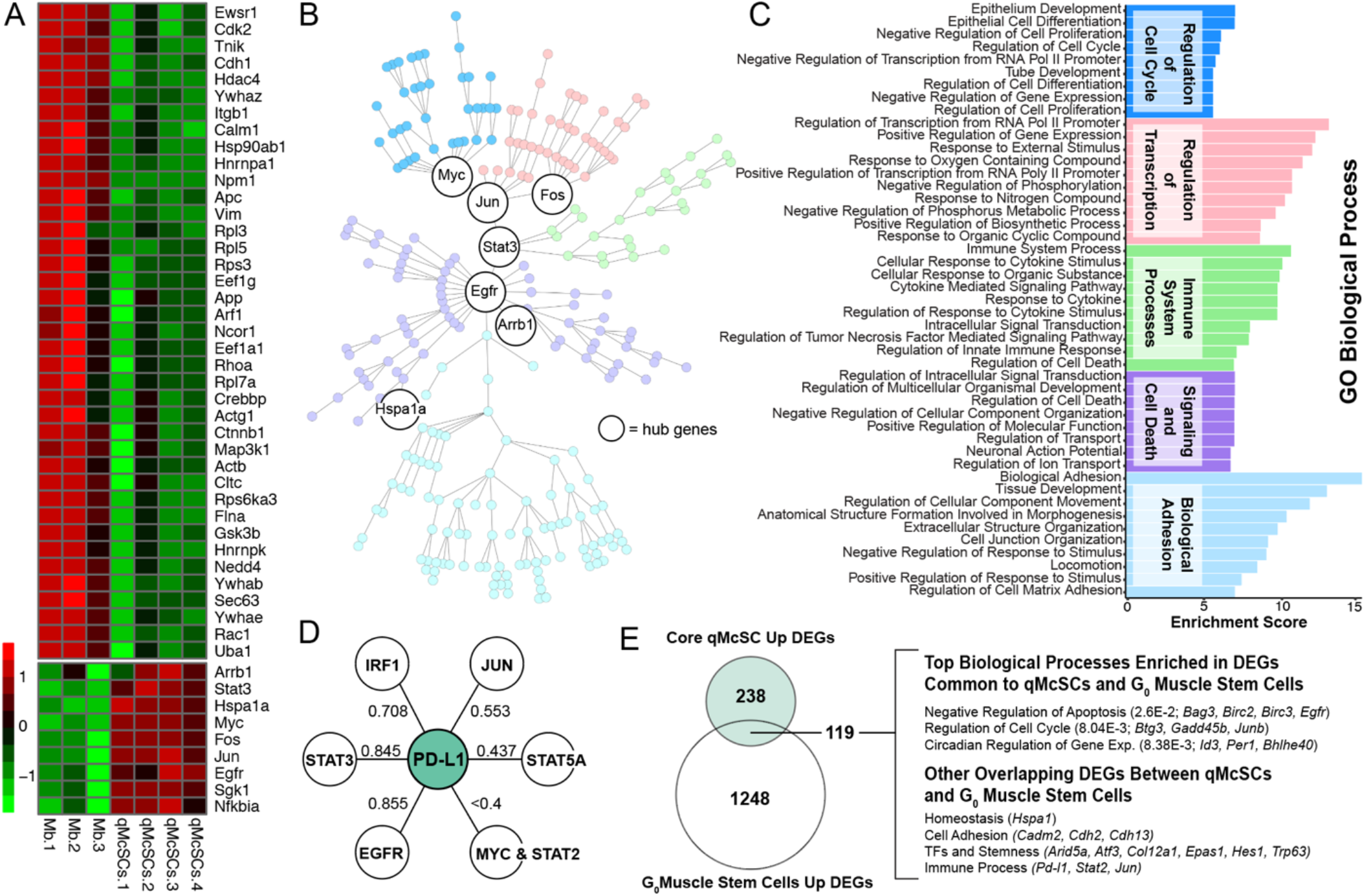
A) Heatmap of hub genes differentially expressed by P0.5 melanoblasts and 8-week adult qMcSCs. (B) Protein-protein interaction network of the upregulated qMcSC core DEGs revealing five major branches enriched for specific biological processes. (C) Clustered bar graph of the top 10 enriched biological processes for each of the five branches identified in B. (D) Combined interaction scores between PD-L1 and upregulated qMcSC core DEGs that are known *Pd-l1* gene promoter binding partners or upstream of *Pd-l1* expression. Interaction scores were based on text-mining, experimental, and database evidence obtained from STRING database (v11.0). (E) Overlap of adult qMcSCs core DEGs and genes with elevated expression in G_0_ muscle stem cells showing a high degree of overlap (119/357). Select biological processes associated with these common genes are indicated.

To further delineate the biological pathways upregulated by qMcSCs, we used functional connectivity to highlight the upregulation of hub genes and their downstream targets within our data. Interconnectivity between the proteins of the 357 genes upregulated in qMcSCs was determined using the STRING (v11.0) database and the network was generated using the Cytoscape (v3.7.1) software package (Szklarczyk et al., 2019; Shannon et al., 2003; Su et al., 2014). The resulting interactome consisted of 265/357 genes with known or predicted PPIs (confidence score > 0.4; text mining, experimental, and database) underlying the regulation of qMcSCs (**Figure 2b**). To uncover downstream biological processes regulated by the previously identified gene hubs, the network was optimized to a radial layout after applying the Kruskal maximal spanning algorithm to trim the total number of edges leaving only the highest combined evidence scored associations between nodes, which resulted in five major branches. Enrichment analysis of the five branches revealed that each includes major biological phenomena that we generally describe as regulation of transcription, biological adhesion, regulation of cell cycle, signaling and cell death, and immune system processes (**Figure 2c**). This finding highlights processes with known roles in G_0_ regulation like biological adhesion and regulation of cell cycle, but again highlights the novel process of the immune system as an integral component of the regulation of the qMcSC state.

Further investigation of the normalized RNAseq read counts revealed the immune checkpoint inhibitor *Cd274* is expressed within both melanoblasts and qMcSCs yet is significantly upregulated in the latter. *Cd274* (referred to here as gene name *Pd-l1)* encodes the protein program death-ligand 1 (PD-L1) and was of particular interest as PD-L1 is an immune-evasion mechanism that is coopted in melanoma, an aggressive form of skin cancer involving melanocytes (Dong et al., 1999; Freeman et al., 2000). PD-L1 inhibitors used in cancer immunotherapy were first approved for patients with this disease and have since expanded to include several cancer sub-types (Akinleye and Rasool, 2019; Raedler, 2015). In addition to *Pd-l1* upregulation in qMcSCs, we also found increased expression in *Egfr*, a signal receptor upstream of *Pd-l1* transcriptional activation (Zhang et al., 2016), and several genes whose proteins are known to directly bind or regulate *Pd-l1* including the hub genes *Stat3, Jun* and *Myc* along with the transcription factors *Irf1, Stat5a* and *Stat2* (Casey et al., 2016; Garcia-Diaz et al., 2017; Yi et al., 2021). The combined interaction scores between PD-L1 and EGFR, STAT3 and IRF1 (determined by STRINGv11) were found to be the strongest at 0.855, 0.845, and 0.708, respectively (**Figure 2d**). This observation highlights a potential regulatory mechanism involving EGFR, STAT3, and IRF1 in driving PD-L1 expression during G_0_ induction in McSCs. These observations are notable because a mechanistic role for PD-L1 in the skin has been insinuated. A small study showed that intradermal injection of anti-PD-L1 inhibitors during early anagen resulted in accelerated hair growth (Zhou et al., 2021). Evidence suggesting a role for PD-L1 in qMcSC regulation also exists. Clinical reports showing elderly patients treated for lung cancer or lymphoma with immunotherapy targeting PD-1/PD-L1 signaling experienced a dramatic side effect— near-complete repigmentation of their age-associated gray hair (Manson et al., 2018; Rivera et al., 2017). Based on these observations we hypothesize that immune system processes play an important role in qMcSCs and that PD-L1 is a marker and potentially a mediator of McSC G0. Further support for increased *Pd-l1* expression during G_0_ was confirmed through secondary analysis of an existing dataset comparing gene expression changes between qMcSC and their proliferating progeny (Infarinato et al., 2020). In agreement with our findings that higher expression of *Pd-l1* is associated with qMcSCs, data from Infarinato et al. shows that the transition from qMcSC to differentiated McSC progeny results in decreased expression of *Pd-l1*.

In order to understand the broader relevance of our results we also asked whether the core-DEGs identified in qMcSCs are common to those in other G_0_ stem cell populations. To answer this, we took advantage of a publicly available gene expression dataset comparing muscle stem cells that reside either in a shallow G_0_ state, known as G_Alert,_ or a fully G_0_ state associated with a reduced rate of reactivation and proliferation (Rodgers et al., 2014). In total, 3241 DEGs (q-value < 0.05, absolute L2FC > 0.5) were found between these two G_0_ states. Regardless of gene expression change, a considerable number of genes overlapped with the qMcSC core-DEGs identified above (133/357 genes, 37.5%). Notably, those genes upregulated in the deeper G_0_ state in muscle stem cells represented the majority of overlapping expression between these two G_0_ stem cell populations (119/133). Analysis of these overlapping genes showed biological processes associated with negative regulation of apoptosis (*Bag3, Birc2, Egfr*, and *Birc3*), regulation of cell cycle (*Btg2, Gadd45b*, and *Junb*) and circadian regulation of gene expression (*Id3, Per1*, and *Bhlhe40*) to be enriched in both G_0_ melanocyte and muscle stem cells (**Figure 2e)**. This overlap also included genes related to protein homeostasis (*Hspa1*), adhesion (*Cadm2, Cdh2*, and *Cdh13*), TFs, and stemness (*Arid5a, Atf3, Col12a1, Epas1, Hes1*, and *Trp63*) that we noted above in qMcSCs (Figure 1b). Lastly, *Pd-l1* is among the genes expressed at higher levels in both qMcSCs and the more deeply G_0_ muscle stem cells, along with the TFs that show connectivity to *Pd-l1* (Figure 2d), *Stat2* and *Jun*. These results suggest that G_0_ may be regulated using common mechanisms in melanocyte and muscle stem cell populations and that expression of *Pd-l1* and other immune-related genes may play unappreciated roles in stem cell G_0_ *in vivo*.

### PD-L1 expression at the cell membrane marks a subpopulation of quiescent McSCs *in vivo*

To test whether increased transcript abundance of *Pd-l1* during G_0_ translates into a spatio-temporal expression pattern of PD-L1 protein that matches when McSCs are G_0_ *in vivo*, we evaluated PD-L1 expression across distinct hair cycle stages. Specifically, skin from 2-4-month-old male and female mice was harvested during dormant (telogen), active (anagen), and regressive (catagen) stages of the hair cycle and processed for immunolabeling. During telogen, PD-L1 is expressed robustly by the innermost layer of hair follicle bulge cells in a position overlapping the location of keratin 6 (K6)-expressing hair follicle cells, and more faintly by the surrounding, outer root sheath layer of hair follicle stem cells and the secondary hair germ. During anagen, PD-L1 expression is modest to absent in the companion layer of the upper hair follicle but not observed in the hair bulb. During catagen, PD-L1 is upregulated in a staining pattern similar to that of telogen with PD-L1 also present in the diminishing epithelial strand in the regressing portion of the hair follicle. PD-L1 is also detectable in the basal layer of the epidermis. Using the melanocyte lineage marker DCT, we clearly show the McSC population localized to the upper permanent portion of the hair follicle during all three stages of the hair cycle **(Figure 3a)**. At telogen, in particular, McSCs reside within the outer layer of the hair follicle bulge (not shown) as well as in the secondary hair germ and some appear to also exhibit faint PD-L1 immunolabeling at the cell membrane. Melanocytes in the hair bulb of anagen stage hairs do not exhibit PD-L1 expression.

**Figure 3.**
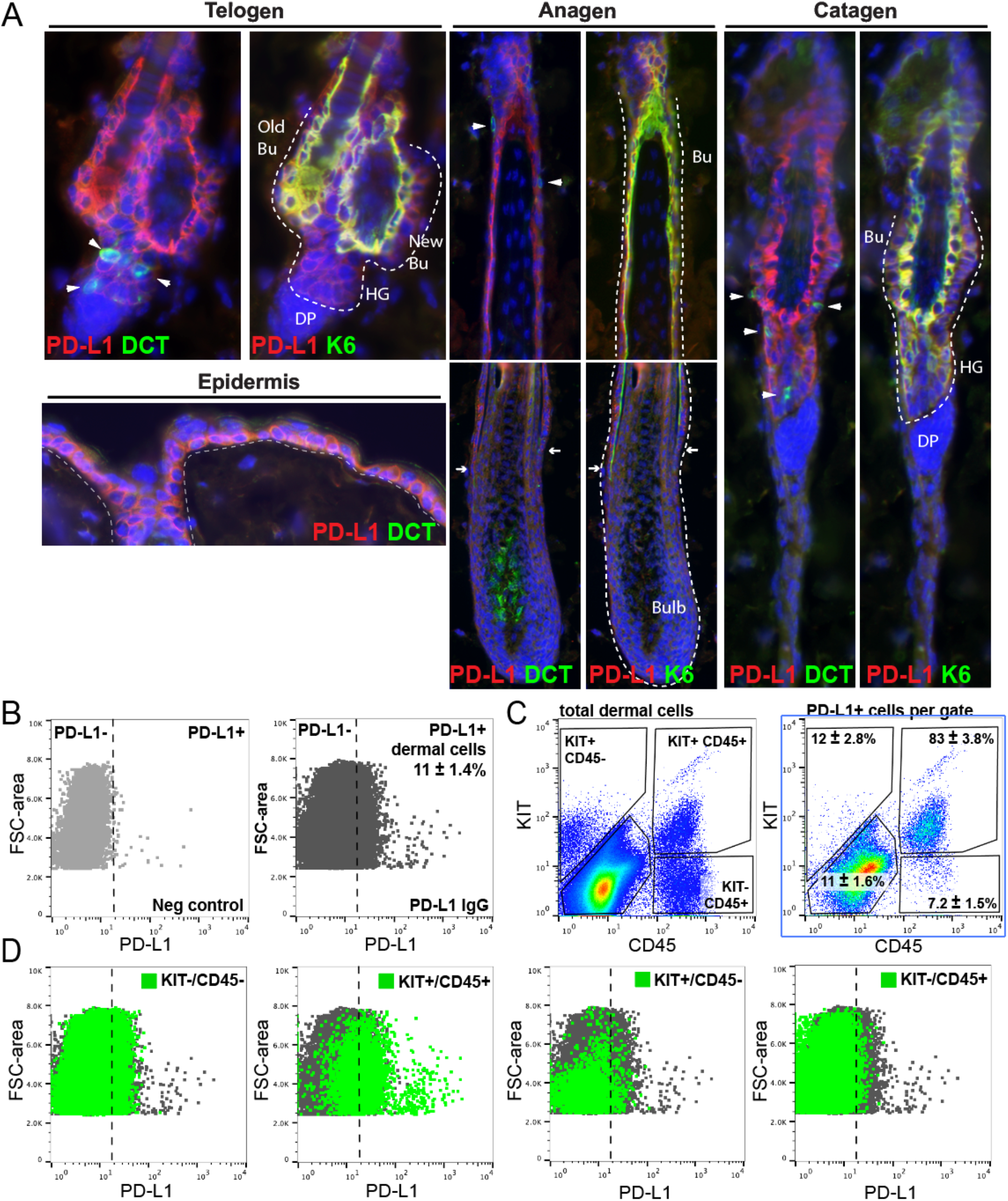
(A) Representative images of PD-L1 staining (red) of the hair follicle and McSCs (DCT, green) during the and telogen (dormant), anagen (active), and catagen (regression) stages of the hair cycle and epidermis in 2-4-month old males and females. The K6+ (green) inner layer of hair follicle cells (at telogen and catagen) and the companion layer (at anagen) is indicated matched images. Tissues were counterstained with DAPI (blue). Arrowheads mark McSCs. Arrows in the anagen images indicate the extent of PD-L1 staining. Dotted lines highlight the shape of the hair follicles. Bu, bulge; HG, hair germ. (B) Gating strategy for flow cytometric assessment of PD-L1^mem+^ cells within the dermis of telogen skin. (C) Gating strategy for the quantification of qMcSCs (KIT+/CD45-), mast cells (KIT+/CD45+), and other immune cells (KIT-/CD45+) along with percent PD-L1^mem+^ staining of these subpopulations (n=4). (D) PD-L1 expression intensity and cell size (FSC-are) across the various KIT/CD45 populations of the dermis.

Using the more sensitive and quantitative technique of flow cytometry, the expression of PD-L1 by McSCs during telogen was confirmed. Dermis from 8-week-old, adult mice were disassociated and assessed for PD-L1 membrane expression, which is detectable in non-permeabilized cells and referred to here as PD-L1^mem+^ with non-stained cells referred to as PD-L1^mem-neg^. On average, based on gating above the background of non-stained cells, 11± 1.4% of the total dissociated dermal cells were PD-L1^mem+^ (**Figure 3b**). As suspected, a portion of KIT+/CD45-McSCs expressed PD-L1^mem+^ (12± 2.8%), as did the majority of KIT+/CD45+ mast cells (83± 3.8%) and some KIT-/CD45+ cells that are presumed to be other CD45-expressing immune cells (7.2± 1.5%), the latter two groups of which are known to express PD-L1 physiologically (**Figure 3c**)(Hirano et al., 2021; Qin et al., 2019). We anticipate that the remaining KIT-/CD45-cells that are also colabeled with PD-L1^mem+^ (11 ± 1.6%, **Figure 3b**) correspond in part to the PD-L1+ inner layer of hair follicle bulge cells identified in immunolabeled tissue sections (**Figure 3a**). PD-L1 ^mem+^ fluorescence intensity also matches that observed in tissue sections with KIT+/CD45+ mast cells and KIT-/CD45-presumed hair follicle bulge cells having the highest PD-L1^mem+^ intensities, while KIT+/CD45-McSCs and KIT-/CD45+ immune cells generally exhibiting lower intensities (**Figure 3d**). In corroboration of our cell sorting approach, KIT+/CD45-McSCs are relatively small in comparison to the total dermal population (based on forward scatter, a measure of cell size; **Figure 3d**) and is an observation that coincides with previous reports (Nishimura et al., 2002)).

Only recently has PD-L1 expression within hair follicles been interrogated (Zhou et al., 2021). Our immunohistochemistry confirms these previous reports that PD-L1 expression fluctuates with hair cycling but extends these initial observations to demonstrate PD-L1 expression at both catagen and telogen stages and more definitely localizing high and low PD-L1 expression to distinct layers of the hair follicle and some McSCs. Additionally, flow cytometric analysis further confirms the novel discovery that PD-L1 is not only expressed at the cell membrane of immune cells within the dermis at telogen but also by some non-immune cells, including a portion of the qMcSC population. Together, this expression pattern advocates for a physiological role for PD-L1 in the hair follicle and McSC dynamics during the regressive and dormant stages of the hair cycle. These results also show that PD-L1^mem+^ can be used to identify a subpopulation of qMcSCs *in vivo* that exist in early adulthood and parallels the upregulation of *Pd-l1* gene expression seen in our qMcSCs core-DEGs.

### Developing a method to quiesce melanoblasts *in vitro* to interrogate changes in PD-L1 expression

To attribute relevance to PD-L1^mem+^ expression by qMcSCs *in vivo*, we next asked whether increased PD-L1 is associated specifically with the G_0_ program (in contrast to a cell cycle-independent stem cell program). To test this possibility, we used the *in vitro* culture method of serum deprivation and mitogen withdrawal to specifically induce G_0_ in the melb-a melanoblast cell line. Melb-a cells are an immortalized, non-differentiated, non-stem cell line derived originally from McSC precursors isolated from neonatal skin (Sviderskaya et al., 1995). For this G_0_ induction method, cells are synchronized to the G_1_ phase of the cell cycle using mitogen withdrawal (removal of SCF and bFGF) for one day followed by reducing serum from 10% to 0.1% to induce G_0_ and then evaluated using flow cytometry. 5-ethynyl-2’-deoxyuridine (EdU) labeling, which is widely used to visualize cells actively synthesizing DNA (Salic and Mitchison, 2008), demonstrates that two days of serum deprivation (aka, 2-day G_0_ cells) is sufficient to stall melb-a cells in the G_0_/G_1_ stages of the cell cycle. This contrasts with actively proliferating melb-a cells grown in media with mitogens and 10% serum (‘active cells’), which are found distributed across all stages of the cell cycle including the S phase (**Figure 4a**). Immunofluorescent staining showed that 2-day G_0_ cells also downregulate the classic cell cycle and proliferation marker mKI67 with 2-day G_0_ cells exhibiting roughly 50% of the mKI67+ cells observed in active cells (**Figure 4b**). Since G_0_ is defined by reversible arrest, we confirmed the ability of 2-day G_0_ cells to re-enter the cell cycle as confirmation of our G_0_ methods. To achieve this, we reintroduced the required growth factors for proliferation (mitogens plus 10% serum) to 2-day G_0_ cells and used EdU to measure cell cycle reentry. By 18 hours post-activation, reactivated 2-day G_0_ cells are distributed across all phases of the cell cycle, with a similar percentage of cells in the S- and G2/M-phase of the cell cycle when comparing reactivated 2-day G_0_ (45.6%) to active cells (50.6%). Further analysis showed, that in reactivated 2-day G_0_ cells the majority of EdU+ cells are concentrated in the S-phase of the cell cycle, a clear indication that G_0_ melb-a cells quiesced using our conditions can successfully reenter the cell cycle. **(Figure 4c)**. Taken together, these results support the use of this *in vitro* mitogen withdrawal plus serum deprivation method to evaluate the nature of G_0_ using the melb-a cell line.

**Figure 4.**
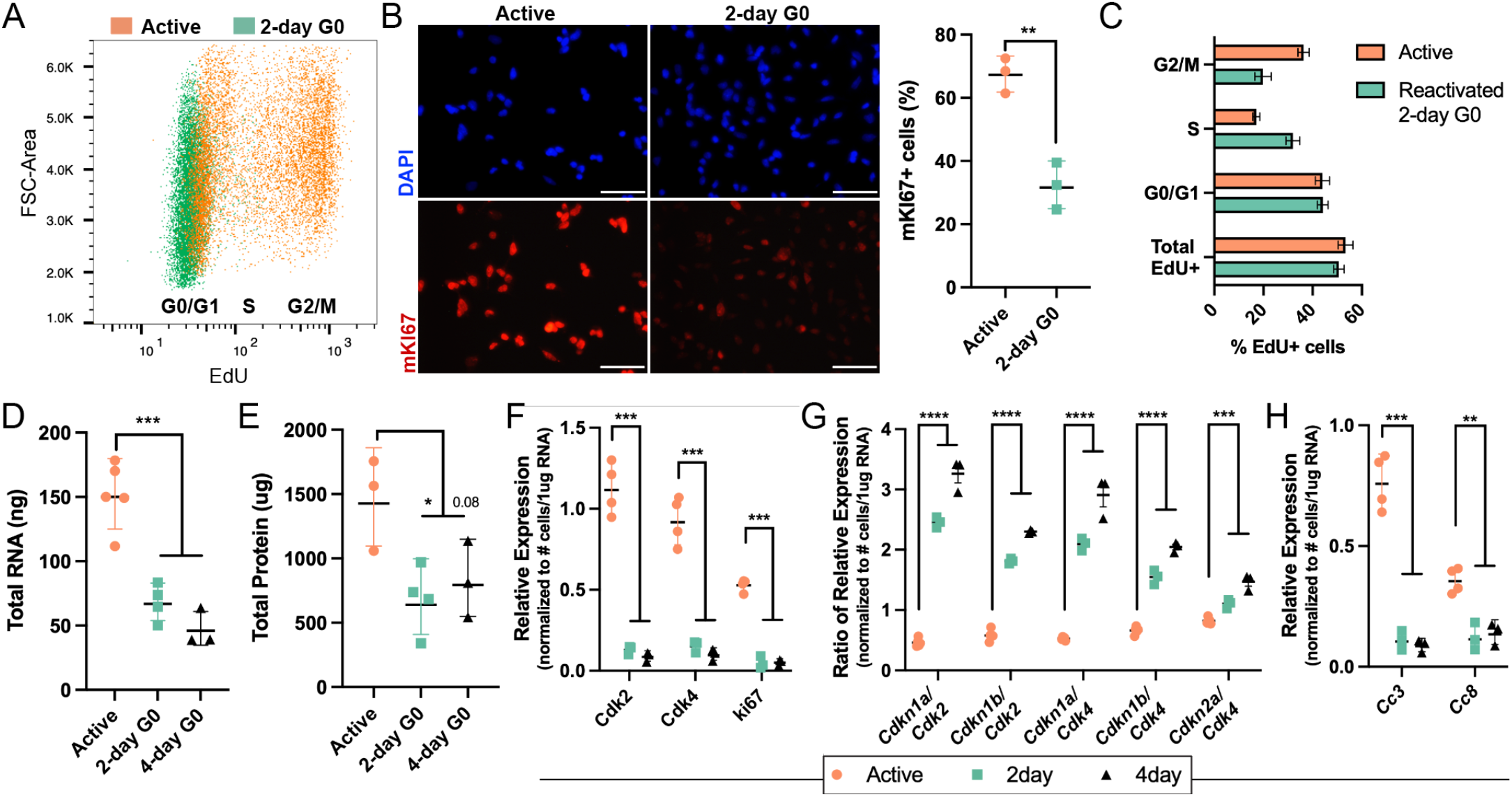
(A) EdU labeling of actively proliferating and 2-day G_0_ melb-a cells demonstrating that 2-day G_0_ are not actively synthesizing DNA and have exited the cell cycle. (B) Representative images of actively proliferating and 2-day G_0_ melb-a cells stained for DAPI (blue) and mKi67 (red), and associated quantification of % mKi67+ cells assessed by flow cytometry. (C) Incorporation of EdU 18 hours post reactivation of 2-day G_0_ melb-a cells compared to actively proliferating cells confirms the ability of 2-day G_0_ cells to reenter all stages of the cell cycle after quiescence induction. (D-E) Quantification of total RNA and protein present in 1×10^6^ cells actively proliferating 2-day G_0_, and 4-day G_0_ melb-a cells demonstrating a reduction in overall RNA and protein levels. (F) RT-qPCR quantification showing decreased relative expression of proliferation markers (*Cdk2, Cdk4*, m*Ki67*) in 2-day G_0_ and 4-day G_0_ compared to actively proliferating melb-a cells. (G) RT-qPCR quantification represented by a ratio of cyclin-dependent kinase inhibitors to respective CDKs demonstrating an increase in 2-day G_0_ and further increased in 4-day G_0_ compared to actively proliferating melb-a cells. (H) RT-qPCR quantification showing decreased relative expression of apoptosis markers (*Cc3* and *Cc8*) in 2-day G_0_ and 4-day G_0_ compared to actively proliferating melb-a cells. In column graphs, each dot represents a biological replicate and bars represent the mean ± SD. Asterisks indicate adjusted p-value (* < 0.05, ** < 0.01, *** < 0.001, **** < 0.0001).

Generally described as a static state of inactivity, cells suspended in G_0_ show downregulation of several molecular processes including DNA, RNA, and protein synthesis (Augenlicht and Baserga, 1974; Gray et al., 2004). In comparison to active melb-a cells, 2-day G_0_ cells exhibit significant reductions in their total RNA and protein levels (as measured by spectrophotometer) with actively proliferating cells having roughly two times the total RNA and protein content compared to G_0_ cells. This low level of RNA and protein remains relatively consistent even if G_0_ is extended to 4 days, suggesting that the G_0_ program is likely established within the initial 2-day timeframe (**Figure 4d-e**). We further characterized this G_0_ program by evaluating the expression of cell cycle genes using qPCR. Two major assumptions are generally made with regards to qPCR experiments: (i) that only a small number of genes are changed between the two sample conditions, and (ii) that the overall size of the transcriptome (RNA content) remains constant across sample conditions (Lovén et al., 2012). We have shown this is not the case for both assumptions. To address these issues, we normalized the mean gene expression values by cell number rather than a standard housekeeping gene; the total cell number required to generate 1ug of RNA was determined by cell sorting. Using this method, 2-day, and 4-day G_0_ cells exhibit significant reductions in the expression of the early cell cycle activator genes *Cdk2* and *Cdk4*, as well as *Ki67* (**Figure 4f)**. Genes for the known inhibitors of CDKs, *Cdkn1a* (p21), *Cdkn1b* (p27), and *Cdkn2a* (p16), conversely show an increased ratio of expression when compared to their respective interaction partners. As the length of G_0_ was extended we observed a significant increase in the ratio of CDKNs to CDKs in 4-day G_0_ cells compared to 2-day G_0_ cells suggesting that increased lengths of G_0_ might alter the rate at which these cells reenter the cell cycle and proliferate (**Figure 4g**). We also confirmed there was a similar reduction in expression of the apoptosis markers cleaved caspase 3 and 8 (*Cc3, Cc8*) to further rule out the possibility that 2- and 4-day G_0_ cells have simply switched to an apoptotic program (**Figure 4h**). In line with our EdU staining, these results indicate that melb-a cells grown in G_0_ media are indeed quiesced; they exhibit classic markers of G_0_ and have decreased expression of cell cycle genes but retain the ability to reenter the cell cycle when appropriate conditions are present.

### PD-L1 is a biomarker for quiescence in melanoblasts *in vitro*

Having established the G_0_ nature of melb-a cells after growth in G_0_ media and confirming their ability to reactivate and re-enter the cell cycle, we next asked whether these cells also upregulate *Pd-l1* RNA and PD-L1 protein similar to that observed in qMcSCs *in vivo*. Akin to our RNA-seq results we find that *Pd-l1* gene expression is significantly increased in 2- and 4-day G_0_ cells compared to active cells (**Figure 5a**). By immunocytochemistry, PD-L1 protein is clearly expressed by both active and 2-day G_0_ cells yet undergoes a change in subcellular localization from perinuclear to a more diffuse, membrane or cytoplasmic pattern upon G_0_ induction (**Figure 5b**). Using flow cytometry, we confirmed that at least part of this pattern is due to PD-L1 shuttling to the cell membrane of G_0_ cells; 2-day G_0_ cells exhibit a significant increase in the percentage of cells expressing PD-L1^mem+^ as well as a significant increase in the geometric mean fluorescence intensity (gMFI) of PD-L1^mem+^ cells over actively proliferating cells (**Figure 5c**). Interestingly, both the percentage and gMFI of PD-L1^mem+^ cells continue to increase with the length of G_0_ (**Figure 5c**). Given that PD-L1 function and stability relies on its acquisition of post-translational modifications (Hsu et al., 2018; Li et al., 2016), namely *N-*linked glycosylation, we also evaluated for changes in PD-L1 size by western blotting using the same number of cells across the varying cell states tested. Melb-a cells exhibit three distinct bands that label with PD-L1 antibody (∼35, 39, and 45 kDa), a pattern consistent with known PD-L1 *N*-linked glycosylation patterns (Li et al. 2016). The longer the cells are in G_0_, the abundance of protein in the PD-L1 banding pattern shifts towards the highest molecular weight band, presumably a highly glycosylated version of PD-L1 (**Figure 5d**). This change in the PD-L1 banding pattern is in distinct contrast to the decrease in ubiquitous proteins like ACTIN and PCNA, the latter of which corresponds to the reductions in total protein abundance observed with G_0_ cells (**Figure 5d;** compare to Figure 4e). These *in vitro* observations parallel our *in vivo* discovery that *Pd-l1* gene expression is upregulated with G_0_ in McSCs and that a portion of qMcSCs in dormant hairs exhibit PD-L1^mem+^ expression (**Figure 2, 3**). Altogether, these data provide further plausibility that PD-L1 is in the right place (cell membrane), in the right state (likely glycosylated) and at the right time to play a functional role during G_0_ in cells of the melanocyte lineage.

**Figure 5.**
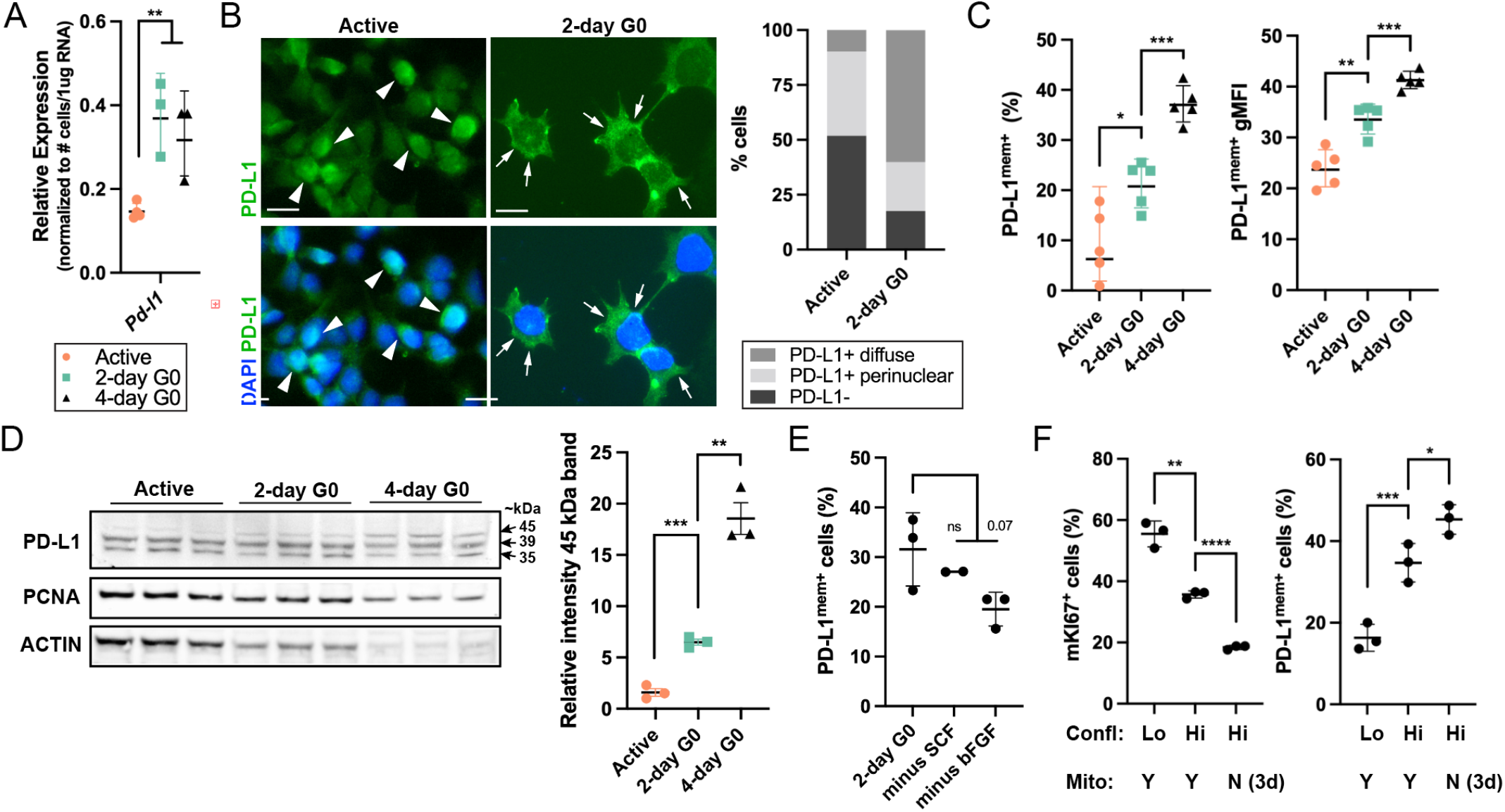
(A) RT-qPCR quantification showing increased relative expression of *Pd-l1* (*Cd274* gene) following G_0_ induction in 2- and 4-day G_0_ melb-a cells. (B) Representative images of actively proliferating and 2-day G_0_ melb-a cells stained for DAPI (blue) and PD-L1 (green) demonstrating two different PD-L1 expression patterns, perinuclear and diffuse. Bar graph represents the associated quantification of these staining patterns revealing that 2-day G_0_ melb-a cells exhibit the more diffuse PD-L1 staining pattern. (C) Flow cytometric quantification of PD-L1^mem+^ expression in non-permeablized melb-a cells. Comparison of actively proliferating, 2-day G_0_, and 4-day G_0_ melb-a cells showing an increase in the percentage of PD-L1^mem+^ cells and PD-L1^mem+^ geometric mean fluorescent intensity (gMFI) with length of G_0_. (D) Western blot of total PD-L1, PCNA and ACTIN in cell lysates harvested from 1×10^6^ melb-a cells. Bar graph represents quantification of the relative intensity of the 45 kDa PD-L1 band. (E) Single mitogen removal during entry into G_0_ shows that SCF, but not bFGF, is sufficient to induce a similar percentage of PD-L1^mem+^ melb-a cells as compared to 2-day G_0_ melb-a cells. (F) The use of high confluency as an alternative method for G0 shows a similar trend in the decrease of mKi67+ melb-a cells and an in increase in PD-L1^mem+^ melb-a cells but not to the same extent compared to mitogen withdrawal. In column graphs, each dot represents a biological replicate and bars represent mean ± SD. Asterisks indicate adjusted p-value (* < 0.05, ** < 0.01, *** < 0.001, **** < 0.0001).

A number of established morphological and molecular characteristics demonstrate that G_0_ regulation differs based on the type of cells being studied and the conditions used to induce G_0_ (Cheung and Rando, 2013; Coller et al., 2006; Rumman et al., 2015). For instance, our *in vitro* method for inducing G_0_ in melb-a cells is achieved by withdrawing the mitogens SCF and bFGF followed by serum deprivation (aka, 2- or 4-day G0). During hair cycling *in vivo*, SCF is diminished during telogen and highest around the proliferating hair bulb during anagen (Mak et al., 2006; Peters et al., 2003). This suggests that loss of SCF may be a key factor in regulating G_0_ in this context, and we were curious to know whether our *in vitro* model reflects this requirement. Indeed, eliminating just one mitogen at a time during G_0_ induction *in vitro* indicated that withdrawal of SCF, but not bFGF, is sufficient to induce a similar percentage of PD-L1^mem+^ cells as with full mitogen withdrawal (**Figure 5e**). In addition to mitogen withdrawal, we also found that induction of G_0_ using contact inhibition another common method for inducing G0 (cells grown at high confluency in 0.1% serum and in the presence of SCF and bFGF) resulted in a similar (but less robust) inverse relationship between PD-L1^mem+^ and Ki67 expression as observed with mitogen withdrawal (**Figure 5f**). Altogether these observations indicate that endogenous PD-L1^mem+^ expression is associated with G_0_ in the melanocyte lineage both *in vivo* and *in vitro* and may rely predominately on SCF removal. Our *in vitro* work also demonstrates that the change in PD-L1 expression observed upon G_0_ induction is not unique to the method by which it is initiated, but rather, is a feature of G_0_ in these cells. Most importantly, these data counterbalance the abundance of research attributing a role to PD-L1 in cancer and highlight the strong possibility that PD-L1 also participates during G_0_ in a physiological and non-tumorigenic sense.

### The aged qMcSC pool is overall more quiescent and comprised of a larger portion of PD-L1^mem+^ qMcSCs

Depletion of the McSC pool with age is considered the primary cause of gray hair (Nishimura et al., 2005), however, the phenomenon of gray hair reversal suggests not all McSCs are lost with age (Yale et al., 2020). Beginning at 12 months of age, the maximal length of the telogen stage of the hair cycle gradually increases by up to 32% during a process known as telogen retention (Chen et al., 2014; Cho et al., 2016; Keyes et al., 2013). A consequence of telogen retention is that McSCs are suspended in the G_0_ state for increasing lengths of time with age. Previous studies also suggest that the length of G_0_ can affect G_0_ depth and the ability of G_0_ cells to reenter the cell cycle (Kwon 2017). Based on these ideas, we hypothesize that the qMcSC pool is molecularly distinct in young and aged animals and these molecular changes will reflect a McSC’s potential for reactivation. To specifically address this question at the transcriptional level we compared gene expression changes between qMcSCs isolated from young and aged female mice using similar RNA-seq analysis methods described above. Using gray hair as a surrogate for age-related McSC pathology, we first confirmed that 24 months is a sufficient age to lead to McSC dysfunction. We collected pelage hairs from the lower back region of adult (9-week, n=5) and aged (24-month, n=4) C57BL/6J female mice by depilation and quantified the presence of pigmentation within the hair shafts as previously described (Anderson et al.). In adult mice, an average of 90% of hairs exhibited high to medium levels of pigmentation whereas in aged mice an average of 65% of hairs show low to no pigmentation with the overall frequency of fully nonpigmented, white hairs reaching 15% (**Figure 6a**). Based on this graying phenotype we anticipate this aged timepoint is sufficient to reveal age and graying-related changes in qMcSC transcriptomics.

**Figure 6.**
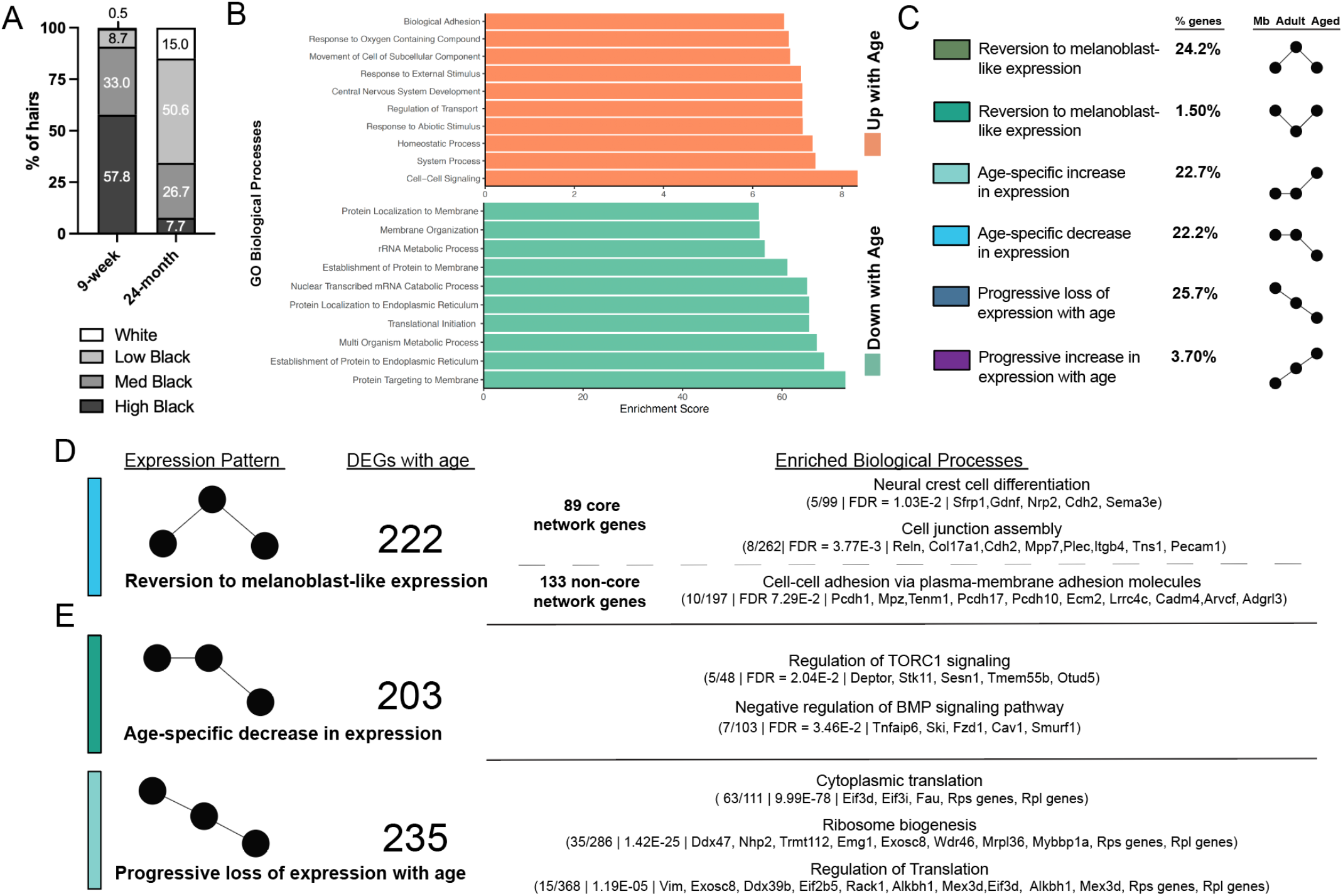
(A) Quantification of the decline in hair follicle pigmentation in 24-month-old aged (n-4) compared to 8-week-old adult female mice (n=5). (B) Clustered bar graph of the biological processes identified by gene set enrichment analysis of the DEGs associated with aged qMcSCs compared to adult qMcSCs. (C) Overview of the expression patterns of the DEGs identified in aged qMcSCs across the melanoblast (P0.5), adult (8-week), and aged (24-month) timepoints. (D-E) Breakdown of the number of DEGs and representative biological processes associated with the expression patterns indicated.

Comparing qMcSCs isolated from adult (8-week-old) and aged (24-month-old) female mice using RNAseq we find 916 DEGs (q-value < 0.05, L2FC > 0.5). Basic gene ontology analysis of the upregulated DEGs (256/916) showed low enrichment for a few broad cellular pathways (e.g., central nervous system development, response to external stimulus, cell adhesion) whereas the downregulated DEGs (660/916) showed high enrichment for a set of related processes including translation, ribosome biogenesis and protein trafficking (e.g., protein targeting to membrane, translational initiation, rRNA metabolic process) **(Figure 6b)**. A fine balance between low translational activity while also maintaining adequate rRNA synthesis and ribosome biogenesis is an important aspect of inactive stem cells that are poised to reenter the cell cycle (Sharifi et al., 2020). Thus, we interpret the downregulation of genes involved in both protein synthesis and ribosome biogenesis to indicate that the aging of qMcSCs is accompanied by an entrenchment into G_0_ (aka, deep G_0_), a state that likely reduces the reactivatability of this aged stem cell pool. Additionally, we found that the senescence-associated secretory phenotype (SASP), which contributes to age-related tissue dysfunction, was not overly represented and only a few upregulated DEGs in aged qMcSCs overlapped with senescence gene lists (comparison made to gene set REACTOME_SENESCENCE_ASSOCIATED_SECRETORY_PHENOTYPE_SASP, www.gsea-msigdb.org; and the SASP Atlas, http://www.saspatlas.com/); these included *Anapc15, Vgf, Serpine1, Krt78, Krt7, Itgax, Igf2, Hgfac* and *C4a*. Notably, other members of the SASP are downregulated including *Fas, Igfbp3, Igfbp7*, and no change was observed in genes encoding other key SASP marker proteins like *Cdkn2a, ll6, Il1, Cxcl8 (*IL8), the *Mmp* genes, the *Cxcl* genes, *Ccl* genes, or the *Timp* genes (Basisty et al., 2020; Coppé et al., 2010). DEGs between adult and aged qMcSCs also contained some overlap with known genes associated with aging (comparison made to the GenAge database); these included the upregulated DEGs *Igf2, Serpine1* and *Sstr3*, and the downregulated DEGs *Fas, Igfbp3, Lepr, Ngfr, Nr3c1, Prkca*, and *Stk11*. A number of these genes participate in the regulation of metabolism, with the upregulation of *Igf2* along with downregulation of *Igfbp3* and *Stk11* pointing toward enhanced mTOR signaling (Johnson, 2018; van Veelen et al., 2011). Altogether, these observations demonstrate that G_0_ in McSCs is not immutable, is uniquely affected by age and this intrinsic aging process is likely SASP-independent. Interestingly, however, the transcriptional changes that accompany aging in qMCSCs point toward two potentially conflicting biological processes, reinforced G_0_ on one hand and activation of mTOR on the other hand.

Taking advantage of our previous RNAseq data (**Figure 1-2**; melanoblasts vs. qMcSCs), we also used a cross-sectional approach to investigate the gene expression changes between the three time points evaluated in this paper: P0.5 (melanoblast), 8-week (adult qMcSC), and 24-month (aged qMcSC). We were particularly interested in addressing whether qMcSCs exhibit any signature indicative of a predisposition for McSC differentiation. With age, in both mice and humans, at least of portion McSCs differentiate prematurely and exhibit ectopic pigmentation within the stem cell compartment. In general, age-related McSC loss and hair graying is attributed to this stem cell differentiation mechanism (Nishimura et al., 2005). Within our cross-sectional data we observed 6 patterns of expression (**Figure 6c);** 236/916 DEGs that reverted to a more melanoblast-like expression level compared to adult qMcSCs, 411/916 DEGs specific to aged qMcSCs, 269/916 DEGs showing either progressive up or down-regulation with age. We focused first on ‘reversion to melanoblast-like expression’ patterns because we anticipated these DEGs might reveal a return to a melanoblast-level of melanocyte differentiation markers that were lost when these cells acquired stemness. However, of the 14 DEGs that were higher in both Mbs and aged qMcSCs, none were pigmentation related and not enriched for any pathway. For DEGs with low expression in both Mbs and aged qMcSCs, we interpret these genes as important drivers of the G_0_ and stem cell state in McSCs that are lost with age. These DEGs can also be subdivided into genes that are upregulated in adult qMcSCs as part of our core G_0_ network (89/222) or non-core G_0_ network genes (133/222) and are enriched for genes highlighting early neural crest cell differentiation and mechanisms for cell-cell adhesion (**Figure 6d**). Stem cell niche structure is an important aspect of stem cell self-renewal and inclusion of the cell adhesion molecule COL17A1 in this category suggests potential defects in G_0_ entry in aged McSCs (Chen et al., 2013b). COL17a1 is hemidesomosal protein essential for maintenance of the hair follicle stem cell niche and loss of *Col17a1* results in reduced TGFβ signaling, the latter of which is necessary to maintain McSC immaturity and G_0_ (Nishimura et al., 2010; Tanimura et al., 2011). The DEG *Sfrp1* is also notable because stabilization of nuclear β-catenin drives McSC proliferation and Mc progeny differentiation and SFRP1 negatively regulates canonical β-catenin activity within the hair follicle through WNT signaling (Hawkshaw et al., 2018; Rabbani et al., 2011). One DEG exhibiting the ‘reversion to melanoblast-like expression’ pattern, but not included in any of these enrichment gene lists, is *Nfatc1*; *Nfatc1* is low in both melanoblasts and aged qMcSCs. Within the hair, *Nfatc1* promotes hair follicle stem cell G_0_ and is inappropriately maintained in aged hair follicles (Horsley et al., 2008; Keyes et al., 2013).It is important to mention that some of these genes are not considered McSC-specific and underscores the possibility that our sorting protocol, while enriched for Mbs or McSCs, fortuitously contains some hair follicle stem cells and provides a unique readout of both cell types simultaneously. Together the DEGs within the ‘reversion to melanoblast-like expression’ pattern point to dysregulation of pathways involved in McSC self-renewal and G_0_, but not immediate McSC differentiation, as contributors to aging within qMcSC.

Of the remaining patterns, ‘age-specific loss of expression’ and ‘progressive loss of expression’ highlight a dichotomy with our gene expression data. Within the category ‘age-specific loss of expression’ we enrich for genes that further support the idea that aged qMcSCs may be less G_0_ and more prone to loss of self-renewal compared to their more youthful qMcSC counterparts. First, aged McSCs have decreased levels of mTOR inhibitors *Deptor, Stk11*, and *Sesn;* mTOR activity defines different states of G_0_; higher levels are associated with more ‘alert’ G_0_ stem cells that can respond rapidly for tissue repair while lower levels are associated with ‘deep’ G_0_ stem cells that are kept in reserve (Rodgers et al., 2014). Second, aged McSCs also have decreased levels of the BMP signaling inhibitors *Ski, Cav1*, and *Smurf1*; during pigment regeneration, BMPs do not appear necessary for maintenance of qMcSCs yet are essential for proper maturation of committed McSC progeny (Infarinato et al., 2020). On the other hand, within the category ‘progressive loss of expression’ we observe enrichment for the translation and ribosome biogenesis pathways identified in Figure 6B that dominate the enrichment signature for DEGs observed in aged qMcSCs. Including melanoblasts into this comparison reveals that downregulation of protein synthesis is not unique to aged qMcSC but is instead a further reinforcement of the G_0_ process acquired during the initial establishment of qMcSCs. Functional heterogeneity of the McSC pool has been observed previously and could explain these conflicting observations (Joshi et al., 2019; Ueno et al., 2014). Due to the nature of bulk RNAseq we are unable to address this possibility transcriptomically, but we anticipate that our data may reflect two independent populations of qMcSCs with different potentials.

To test this idea directly and using PD-L1 ^mem+^ as a biomarker for G_0_, we compared McSCs harvested from the dermis of adult (8-week) and aged (24-month) mice using flow cytometry. Consistent with the canonical stem cell loss pathway, which is likely driven by loss of McSC self-renewal and premature differentiation upon activation, we observed an average of 49% reduction in the overall size of the qMcSC population in aged mice. However, within the remaining McSC population, PD-L1^mem+^ qMcSCs were retained and thus represented a greater proportion of the qMcSC pool than observed in the aged qMcSC pool (**Figure 7a**). Similarly, PD-L1 in telogen-stage hairs of aged mice continues to be robustly expressed (**Figure 7b**).

**Figure 7.**
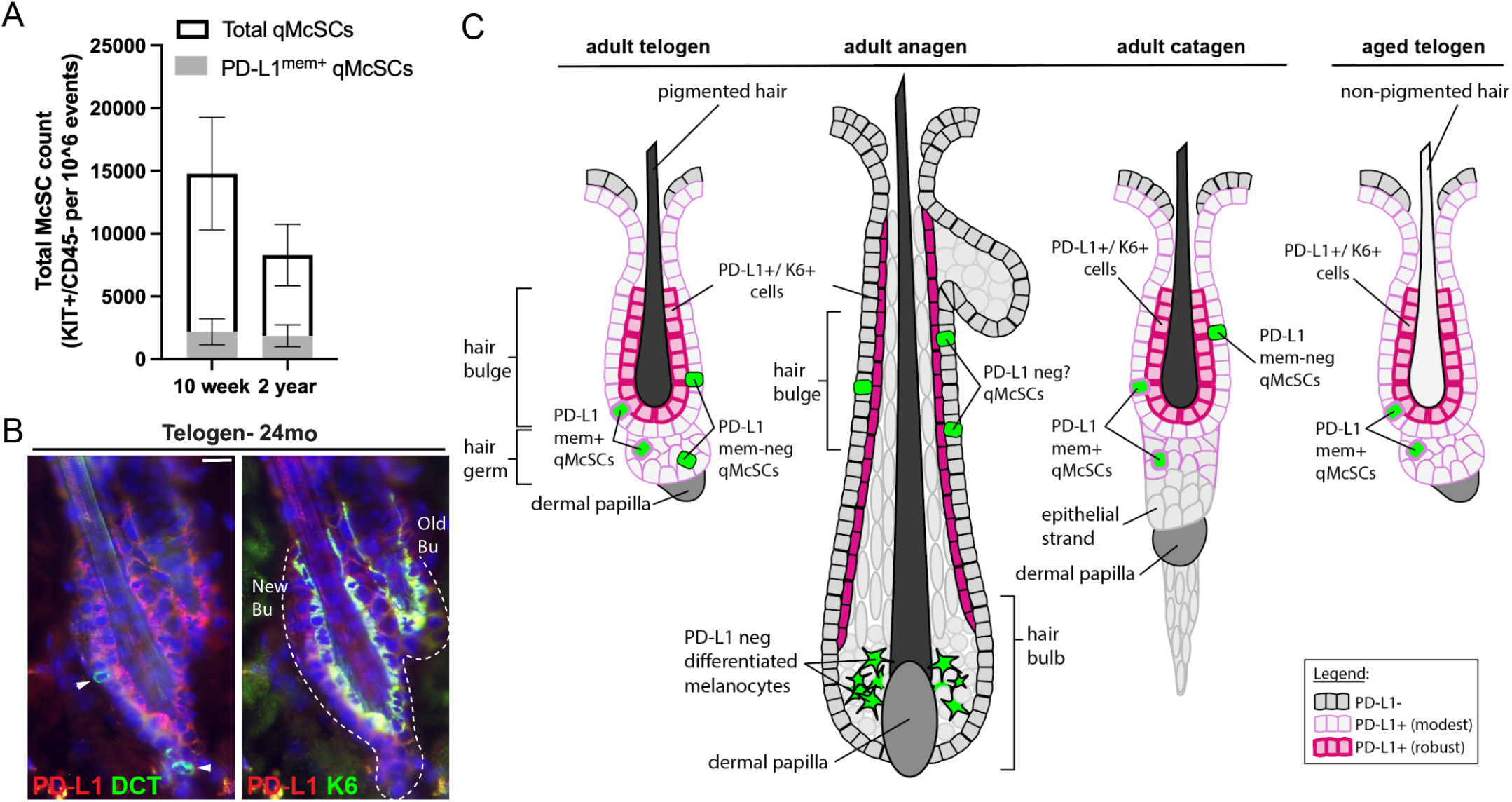
(A) Quantification of PD-L1^mem+^ qMcSCs in adult qMcSCs (10-week) compared to aged (24-month) qMcSCs as determined by flow cytometry of dissociated dermis from female mice (n=3/timepoint). Total qMcSCs are notable reduced with aging whereas the PD-L1^mem+^ subpopulation remains constant. (B) Representative images of PD-L1 (red) and K6 (green) staining of the hair follicle and McSCs (DCT, green) during telogen in aged (24-month) female skin. Tissues were counterstained with DAPI (blue). Arrowheads mark McSCs. Dotted lines highlight the shape of the hair follicles. Bu, bulge; HG, hair germ. (C) Diagram representing the expression of PD-L1 in the hair follicle and McSCs in the context of hair cycling and with age.

The increase in the proportion of PD-L1^mem+^ cells in aged qMcSCs is not predicted by gene expression as *Pd-l1* is not a DEG when comparing adult and aged qMcSCs by RNAseq. A similar observation is made *in vitro*; increasing the duration of G_0_ in melb-A does not alter *Pd-l1* expression at the transcript level (**Figure 5a**) but does lead to changes in PD-L1 protein localization, PD-L1 protein modifications, and a higher proportion of cells exhibiting PD-L1^mem+^ expression (**Figure 5b-d**). This suggests that the post-translational regulation of PD-L1 may be an important aspect of the role of this protein in G_0_ cells. Altogether, these data indicate that the adult qMcSC pool is heterogenous and can be split into two groups defined by PD-L1^mem+^ and PD-L1^mem-neg^ expression. We also show that PD-L1^mem+^ and PD-L1^mem-neg^ subpopulations are affected differently with age and the composition of the qMcSC pool becomes increasingly represented by PD-L1^mem+^ qMcSCs.

## Discussion

In this study, we interrogated the melanocyte lineage at the transcriptomic level to better understand how aging influences G_0_ in McSCs. In contrast to the prevailing view that G_0_ cells are simply transcriptomically depressed versions of their active counterparts, we discovered that adult qMcSCs (8-week), in comparison to actively proliferating melanoblasts (P0.5), upregulate a core network of genes represented by five major biological pathways. We anticipate that the genes in these pathways promote, maintain, and/or protect G_0_ stem cells. One branch of genes within this core network highlighted immune system processes, including upregulation of the immune checkpoint inhibitor *Pd-l1*. In light of the fact that one side-effect of anti-PD-L1 immunotherapy is gray hair reversal (or reactivation of a defunct regenerative pigment system) (Manson et al., 2018; Rivera et al., 2017) our data implicate PD-L1 as a potential driver for repressing qMcSC activation within gray hairs during aging. By employing a novel method to quiesce immortal melanoblasts *in vitro*, we demonstrate that *Pd-l1* transcript and PD-L1 protein expression are both upregulated during the dormant state of G_0_ and that G_0_ induces localization of PD-L1 to the cell membrane where it is known to function in cell signaling. *In vivo*, PD-L1^mem+^ expression identifies a sub-population of qMcSCs as well as K6+ hair follicle cells during the telogen stage of the hair cycle, establishing this protein as a biomarker associated with dormancy (see summary, **Figure 7c**). Notably, PD-L1^mem+^ qMcSCs are retained with age whereas PD-L1 ^mem-neg^ qMcSCs are depleted suggesting that PD-L1 L1^mem+^ qMcSCs may exist in non-pigmented, gray hairs and could be the target of anti-PD-L1 immunotherapy that leads to gray hair reversal (see summary, **Figure 7c**). We anticipate that the aging mechanisms that explain the loss of PD-L1 ^mem-neg^ qMcSCs and retention of PD-L1^mem+^ are reflected in the qMcSC transcriptomics, which suggests two possible biological outcomes. We predict that aged PD-L1^mem-neg^ qMcSCs have reduced self-renewal capacity and will prematurely differentiate upon activation leading to their eventual loss (the canonical pathway for hair graying). Conversely, we predict that aged PD-L1^mem+^ qMcSCs will become increasingly difficult to reactivate as they decline further into G_0_ (a novel, non-canonical pathway for hair graying).

The role of PD-L1 has mostly been explored in the context of immune evasion involving T-cells. Localization of PD-L1 at the cell membrane allows PD-L1 to interact with the programmed death-1 receptor (PD-1) on T-cells, suppressing an immune response and providing cancer cells with immune privilege (Carter et al., 2002; Freeman et al., 2000). In combination with evidence supporting PD-L1 upregulation on cancer cells and its role in escape from the immune response (Iwai et al., 2002; Meng et al., 2018; Zhang et al., 2009), the PD-1/PD-L1 axis has become a significant target for immunotherapy-based clinical cancer research, and treatments targeting this interaction has been approved by the FDA for a variety of cancers including melanoma (Gong et al., 2018). Outside of the cancer context, physiological PD-L1 expression promotes immune tolerance in a variety of human and murine tissues including the thymus, liver, lung, pancreas, eye, and placenta (Bardhan et al., 2016; Keir et al., 2008). Hair follicle immune privilege has long been established and protection of hair follicle stem cells during dormancy is attributed to downregulation of MHC class I molecules that accompanies hair follicle stem cell G_0_ (Agudo et al., 2018; Paus et al., 2003). However, until now, no studies have reported PD-L1 upregulation as a possible driver of this privilege process. PD-L1 expression by some qMcSC and K6+ hair follicle cells at telogen suggest that these cells actively regulate their immune detection status and should be tested directly. A physiological role for PD-L1 in G_0_-mediated McSC immune evasion may also help to explain the increase in vitiligo-like lesions in melanoma patients receiving anti-PD-1 or PD-L1 immunotherapies compared to other immunotherapies (Garrett et al., 2021; Guida et al., 2021).

Outside of immune privilege, evidence directly from cancer research also indicates an association between PD-L1 expression and cell cycle state. IFN-gamma is a popular cytokine used to stimulate membrane expression of PD-L1 in both tumorigenic and non-tumorigenic cells but also causes cell cycle arrest (Chin et al., 1996). However, in our *in vitro* G_0_ model established above, we demonstrate that PD-L1 membrane expression is elevated in melb-a cells by cell cycle arrest alone, independent of IFN-gamma signaling. In astrocytomas, PD-L1^mem+^ expression was highly associated with non-proliferative, KI67-negative cells at the growing edge of tumors (Yao et al., 2009). In this case, PD-L1 expression was independent of tumor stem cell-like properties but dependent on mitogenic availability, and thus the complex microenvironment observed within solid tumors may explain heterogeneity in PD-L1 expression. Others have shown that CDK4 levels, an essential early-stage protein for re-entry into the cell cycle, can influence the abundance of membrane expression of PD-L1 in multiple tumor cell lines. By inhibiting CDK4 expression and causing cell cycle arrest, degradation of PD-L1 is reduced and results in increased PD-L1^mem+^ expression (Zhang et al., 2018). These observations match those that we report in Figure 5b-c demonstrating that differential subcellular localization of PD-L1 (and function of PD-L1) is intimately tied to G_0_ status.

Mechanistically, there is some support for the idea that PD-L1 can influence tissue regenerative properties within the skin. In the context of pigmentation, and as mentioned above, a clinical trial involving PD-1/PD-L1 immunotherapy using function-blocking antibodies patients reported near-complete hair repigmentation in elderly patients (13/14) undergoing treatment for lung cancer (Rivera et al., 2017). These findings suggest that gray hairs maintain the capacity for repigmentation, a fact supported by numerous examples (Yale et al., 2020). Based on our discoveries establishing PD-L1 as a biomarker of G_0_ and the persistence of PD-L1^mem+^ qMcSCs in aged mice, we anticipate that PD-L1 signaling plays a novel role in preserving hyper-dormancy in qMcSCs in gray hairs. Indeed, constitutive expression of PD-L1 in melanoma cell lines results in the decreased expression of several melanin biosynthesis genes including DCT, MITF, MLANA, and TYR a characteristic consistent with G_0_ (Chatterjee et al., 2018). Outside of pigmentation, inhibiting PD-L1 during early entry into the anagen stage of the hair cycle accelerates the hair growth rate further indicating a mechanistic role for PD-L1 in preventing the transition from hair dormancy to hair growth (Zhou et al., 2021).

Several studies have shown that the ability of aged stem cell populations to regulate the transition between G_0_ and proliferation declines with age (Kalamakis et al., 2019; Leeman et al., 2018; Tümpel and Rudolph, 2019). Reduced frequency of stem cell reactivation and tissue regeneration further contributes to the deterioration of tissue integrity and function overtime (Janzen et al., 2006; Krishnamurthy et al., 2006). A deeper state of G_0_ has been used to describe the reduced reactivation rate and prolonged re-entry into the cell cycle of several aged populations of cells *in vivo* including hepatocytes after partial hepatectomy, salivary gland cells after administration of isoproterenol, and muscle stem cell populations following a wounding event (Adelman et al., 1972; Bucher and Swaffield, 1964; Bucher et al., 1964; Liu et al., 2013). However, studies involving parasymbiosis reveal that entrenchment into G_0_ may not necessarily be irreversible as the ability to restore tissue homeostasis is retained in several aged cell populations (hepatic, muscle, and neural) that fail to activate appropriately under normal physiological conditions within an aged environment (Conboy et al., 2003; Iakova et al., 2003; Katsimpardi et al., 2014). *In vitro* studies on rat embryonic fibroblasts may explain why this is the case as holding cells in G_0_ for increasing lengths requires a stronger stimulating signal in order to reactivate at a rate comparable to cells in a shallower G_0_ state (Kwon et al., 2017). These examples in conjunction with the results presented throughout this study point to changes in the depth of G_0_ as a novel paradigm for melanocyte stem cell aging in a subpopulation of qMcSCs. This change in qMcSC attributes may be attributed to the fact that qMcSCs sit for longer and longer periods in G_0_ as the dormant stage of the hair cycle lengthens during the process of telogen retention (Chen et al., 2014; Cho et al., 2016). Future studies focused on better defining the qMcSC pool using single cell RNAseq and functional assays that directly compare the regenerative potential of PD-L1^mem+^ and PD-L1^mem-neg^ qMcSCs are warranted. At a minimum, we anticipate that PD-L1^mem+^ qMcSCs are a viable, in-situ population of deeply quiesced cells that have the potential to be targeted for improved tissue regeneration, which in this case means hair repigmentation. More broadly, evaluating the role of PD-L1 signaling by other G_0_ stem cell populations, like muscle stem cells (**Figure 2e**), is also warranted.

## Acknowledgments

This research was supported by funding provided to MLH from the National Institute on Aging (R00AG047128), the UAB Faculty Development Grant Program, and start-up funds from the Department of Biology and College of Arts of Science at the University of Alabama, Birmingham. Fellowship support was also provided independently to JWP (UAB Blazer Fellowship). We also would like to give special thanks to Zoya T. Anderson and Christopher R. Keys for their valuable insights, feedback, and moral support during this study.

## Author Contribution

Conceptualization: Joseph W. Palmer, Melissa L. Harris

Data curation: Joseph W. Palmer, Kyrene M. Villavicencio, Misgana Idris, Melissa L. Harris

Formal analysis: Joseph W. Palmer, Kyrene M. Villavicencio, Misgana Idris, Melissa L. Harris

Funding acquisition: Joseph W. Palmer, Melissa L. Harris

Investigation: Joseph W. Palmer, Kyrene M. Villavicencio, Dominique Weddle, Misgana Idris, Melissa L. Harris

Methodology: Joseph W. Palmer, Kyrene M. Villavicencio, Misgana Idris, Fabian V. Filipp, NISC Comparative Sequencing Program, Melissa L. Harris

Project administration: Melissa L. Harris, Joseph W. Palmer Resources: William J. Pavan, Melissa L. Harris

Software: Joseph W. Palmer

Supervision: Joseph W. Palmer, Melissa L. Harris

Validation: Joseph W. Palmer, Kyrene M. Villavicencio, Misgana Idris, Melissa Harris Visualization: Joseph W. Palmer, Melissa Harris

Writing – original draft: Joseph W. Palmer, Melissa L. Harris

Writing – review & editing: Joseph W. Palmer, Melissa L. Harris

## Materials and Methods

### Ethics statement

Animal care and experimental animal procedures were performed in accordance with the guidelines set forth by the Public Health Service and Office of Laboratory Animal Welfare as designated by the Institutional Animal Care and Use Committee associated with UAB and NHGRI. This research is associated with the following animal protocols: UAB 20382 (to MLH), NHGRI G-94-7 (to WJP).

### Animals

Animals used for RNA-seq were housed at the animal facilities at NHGRI. C57BL/6J wildtype females (8-week-old) and pups (P0.5) were derived from in-house mating of C57BL/6J mice obtained from The Jackson Laboratory (JAX, JAX:000664, RRID: IMSR_JAX:000664). C57BL/6 wild-type female mice (24-month-old) were obtained through the NIA Aged Rodent Colony (Charles River Laboratories, RRID:SCR_007317) at 21 months old and allowed to age to 24 months within the same facility as the adult 8-week-old and P0.5 pups to which they were compared.

Animals used for flow cytometry, IHC, and hair analysis were housed at the animal facilities at UAB. C57BL/6J wildtype mice used for the flow cytometry (8-week-old female) data presented in Figure 3b-d were obtained from JAX. A combination of C57BL/6 wild-type mice from JAX and the NIA Aged Rodent Colony were used for the IHC data presented in Fig. 3a (2-4-month old males and females) and Fig. 7b (24-month-old females), and the hair shaft color analysis in Fig. 6a. For IHC, some of these animals were depilated before skin harvest to synchronize the hair cycle (anagen and catagen). C57BL/6 wild-type mice (10-week-old and 24-month-old) used for the flow cytometry data presented in Fig.7a were obtained from the NIA Aged Rodent Colony. All animals transferred to UAB were acclimatized within the mouse facilities for at least one week prior to experimentation.

### Melanoblast and McSC enrichment

Dermis from postnatal day 0.5, 8-10-week, and 24-month mouse body skin was dissociated and processed using FACS to obtain enriched populations of melanoblasts, adult and aged qMcSCs, respectively (Harris et al., 2018). Cells of the melanocyte lineage were identified by their surface expression of the receptor KIT. Melanoblasts in the P0.5 dermis are located within developing hair follicles (Peters 2002). Enrichment for quiescent McSCs at both the adult and aged timepoints was achieved by isolating KIT+CD45-cells from mouse skin whose hair was in the telogen stage of the hair cycle. McSC enrichment at telogen is dependent on the fact that telogen hairs contain only McSCs and do not contain differentiated melanocytes (Nishimura 2002). The telogen stage of the hair cycle was confirmed by the pinkness of the skin (Muller-Rover 2001).

In brief, trunk skin was obtained and the subcutis removed by gentle scraping with curved forceps. The remaining skin was incubated at 37°C in 0.25% Trypsin-EDTA (Gibco 25200-056) for 20 minutes and then the dermis was separated from the epidermis by gentle scraping with curved forceps. The resulting dermis was minced using curved surgical scissors and dissociated by enzymatic treatment with 0.3mg/mL Liberase TL (Roche 0540102001) for 45-50 minutes at 37°C. Enzymatic digestion was stopped using a solution of DMEM containing 20% FBS and 0.5 mg/ml Dnase (Sigma DN-25). The dermis was physically disrupted by forcefully passing the dermal cell solution through a 70 µM Filcon filter (070-67S) fitted to a 50mL luerlock syringe (Soft-Ject, 8300006682). Cells were washed using a freshly prepared FACS wash solution consisting of 1X DPBS, 5% FBS, 25mM HEPES (Gibco 15630-080), and 2mM EDTA before being pelleted at 350 x g for 7 min at 4°C. Cells were resuspended and stained using cell surface antibody markers preconjugated to a fluorophore: KIT (CD117, BD Pharmingen 25-1171-82, RRID: AB_469644), CD45.2 (BD Biosciences 553772, RRID:AB_395041) and PD-L1 (Invitrogen 12-5982-82, RRID:AB_466089). Fluorescence was assessed using an S3e Biorad cell sorter equipped with 488/561 lasers or a FACSAria (Becton Dickinson), Cells of the melanocyte lineage were positively selected by gating on the KIT+/CD45.2-cell population while mast cells were negatively selected by removing double-positive KIT+/CD45.2+ cells. Analysis of cell sorting data was performed using FlowJo software (V10.6.0, RRID:SCR_008520).

### RNA isolation

Cellular RNA was purified using the Directzol RNA Miniprep Kit (Zymo R2062). For RNA-seq, RNA was treated with 0.8U/µl RNase inhibitor. RNA was quantified using a Qubit fluorometer. RNA quality was determined by Bioanalyzer and RNA samples with a RIN score >8.5 were used for sequencing. For qPCR, RNA was isolated similarly but quantified using an Epoch spectrophotometer (Biotek).

### RNA-seq and differential gene expression analysis

McSCs from 3-4 animals were pooled to generate enough RNA to serve as one biological replicate. Amplified cDNA was created from 20 ng total RNA using the Ovation RNA-Seq System V2 (Tecan Genomics). The cDNA was fragmented using a Covaris E210 before proceeding to library construction with 1000ng cDNA using TruSeq RNA Sample Prep Kit, version 2 (Illumina) using 10 PCR cycles. Libraries were pooled in equimolar ratio and sequenced together on a HiSeq 2500 with version 3 flow cells and sequencing reagents. At least 60 million 126-base read pairs were generated for each library. Data were processed using RTA 1.18.64 and CASAVA 1.8.2. RNA-seq reads that passed the Illumina platform quality check were used for downstream analyses.

Bam files were assessed for low-quality reads and removed before converting to fastq file format using samtools (v1.9, RRID:SCR_014583) and Bam2fastq (v1.1.0) software. Read quality was assessed using FastQC (v0.11.8, RRID:SCR_014583) and trimming was performed using Trimmomatic (v0.36, RRID:SCR_011848). Reads with a Phred quality score of 30 were pairwise removed and the remaining paired reads were trimmed to an average length of 90 base pairs. Reads were aligned to the Ensembl (RRID:SCR_002344) mouse mm10 reference genome with the STAR (v2.5.2, RRID:SCR_015899) aligner software using default parameters, and count tables were generated using the STAR genecounts feature and full Ensembl GTF annotation file. Differential gene expression analysis was performed using DESeq2 (v1.22.2, RRID:SCR_015687) package in R using default settings. All sequencing data generated in this study are available at NCBI GEO under accession GSE######.

Heatmaps were generated by applying the regularized log transformation (rlog) to the results data frame produced by the DESeq2 package. To generate figures, the rlog values for filtered genes of interest were loaded into the pheatmap (RRID:SCR_016418) R package and scaled by row. Core networks were generated using 0.4 confidence protein-protein interactions using the experimental, database, and text mining information obtained from the STRING (v11.0, RRID:SCR_005223) database. Visualization and optimization of the network layout were performed using Cytoscape (v3.7.1, RRID:SCR_003032) and the plug-in Cyspanning tree (v1.1). In brief, unconnected small trees (2-3 nodes) were removed before applying a maximal Kruskal algorithm on the combined evidence scores for edges and a radial layout was used to visualize the resulting branches. Hub proteins and transcription factors were determined by using publicly available data provided by the Enrichr database (RRID:SCR_001575) and AnimalTFDB (RRID:SCR_001624) respectively. Enrichment analysis of biological processes was performed using the Molecular Signatures Database (MSigDB v6.2, RRID:SCR_016863) through the Gene Set Enrichment Analysis website platform or the Panther Database (Release 17.0, RRID: RRID:SCR_004869). Visualization of data and generation of the clustered bar graphs were performed using the ggplot2 (RRID:SCR_014601) R-package.

### qRT-PCR analysis

For each biological replicate, 1 × 10^6^ cells were isolated by flow cytometry and 1<u>µ</u>g of purified RNA was reverse transcribed into cDNA using the High Capacity cDNA Reverse Transcription Kit (Thermo Fisher Scientific 4368813). A standard curve was used to determine the mean quantity value of target transcripts between conditions and across three technical replicates per sample. TaqMan Fast Universal PCR Master Mix (Thermo Fisher Scientific 4444965) and Taqman gene expression assays were used for amplification. Gene expression assays used in this study included Pdl1 (Cd274, Mm03048248_m1), Cdk2 (Mm00443947_m1), Cdk4(Mm00726334_s1), Cdkn1a (Mm01303209_m1), Cdkn1b (Mm00438168_m1), Cdkn2a (Mm00494449_m1) mKi67 (Mm01278617_m1), Cc3 (Mm01195085), Cc8 (Mm01255716). Expression was assessed on a QuantStudio 3 (Thermo Fisher) using the default fast reaction settings. Gene expression was normalized by cell number to account for the change in the transcriptome size of cells across cellular states.

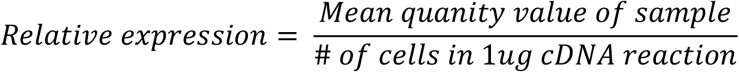

The ratio of cyclin-dependent kinase inhibitors to their respective cyclin-dependent kinases for individual samples across conditions was calculated using the following formula.

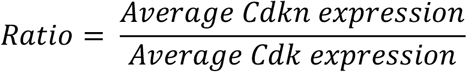

### Immunolabeling

For immunohistochemistry, the skin was immersed in freshly made 2% methanol-free formaldehyde (Thermo Fisher Scientific 28908) in 1X DPBS (w/o Mg and Ca, Gibco 14190-144) for 30 minutes on ice and washed for 24 hours. in 1X DPBS at 4°C. Skin samples were then cryoprotected by placing them in 10% sucrose (Fisher S5-500) in 1X DPBS for additional 24 hours at 4°C prior to freezing in O.C.T. (Fisher 23-730-571). <u>10</u> µm thick cryosections of skin were cut using a cryostat (Thermo Fisher Scientific NX50) and stored at −80. Prior to immunolabeling, slides were immersed into 4% formaldehyde for 15 minutes, washed briefly, and heated to near boiling in 10mM citrate buffer with 0.05% Tween for 2 minutes

For immunocytochemistry, cells were grown on 8- or 4-well chamber slides. Cells were washed with 1X DPBS and fixed using 2% methanol-free formaldehyde in 1X DPBS for 30 minutes at room temperature followed by two rounds of additional washing with 1X DPBS before staining.

Immunolabeling was performed by washing slides in a 1X DPBS and 0.1% Tween 20 (Fisher BP337-500) solution (PBST) for 20 minutes before incubation with primary antibodies overnight at 4°C (for sections) or 30-60 minutes at room temperature (for cells). Slides were washed in PBST solution for 15 minutes followed by incubation with appropriate secondary antibodies (1:2000-1:3000, AlexaFluor, Invitrogen) for 2 hours (for sections) or 30 minutes (for cells). Following antibody incubation, slides were washed in PBST for 15-20 minutes and coverslips mounted with Fluoromount-G with DAPI (Thermo Fisher Scientific 00-4959-52). Primary antibodies used for immunolabeling included: anti-PD-L1 (1:100, Thermo Fisher Scientific 14-5982-82, RRID:AB_467781), anti-DCT (1:400, anti-TRP-2, Santa Cruz Biotechnology sc-10451, RRID:AB_793582), anti-K6a (1:500, Biolegend 905701, RRID:AB_2565052), and anti-KI67 (1:100, Abcam ab16667, RRID:AB_302459). Fluorescence microscopy and imaging were performed either on an EVOS FL Cell Imaging System (Thermo Fisher Scientific) or Eclipse 80i microscope (Nikon). Images were processed using Adobe Photoshop.

### Hair counts

Hair shaft pigmentation was quantified by a binning method developed previously in our lab (Anderson et al.). Briefly, hairs were classified as high, medium, or low black (depending on the amount of white space within the medullary air pockets), and white (essentially no visible pigment granules). All analyses were performed blind to sample age. A minimum of 200 hairs were counted per adult sample (n=5) and over 1000 hairs per aged sample (n=4).

### Cell culture

Melb-a cells (RRID: CVCL_C693) were obtained from the Wellcome Trust Functional Genomics Cell Bank (Sviderskaya et al., 1995). For general cell growth and passaging, cells were incubated at 37°C and 10% CO2 in “full media” consisting of a RPMI 1640 with L-Glutamine (RPMI, Gibco, 11875-093) supplemented with 10% heat-inactivated fetal bovine serum (hi-FBS, Gibco, 16140-071), 100U/mL Penicillin/100ug/mL Streptomycin (pen/strep, Gibco, 15140-122), 20ng/mL SCF (Thermo Fisher Scientific PMC2111) and 0.6ng/mL bFGF (Stemgent 03-0002).

For assessment of actively proliferating cells, cells in sub-confluent T-75 filtered flasks (Thermo Fisher Scientific 156499) were split 1:4 into fresh T-75 flasks, incubated in full media, and harvested 24 hours later at roughly 50% confluency.

For assessment of quiescent cells, G_0_ was induced using serum deprivation or contact inhibition. Serum deprivation was achieved by first syncing subconfluent cells to the G1 stage of the cell cycle by thoroughly washing the cells in 1X DPBS and incubating them in media without the mitogens SCF and bFGF (RPMI, pen/strep, and 10% hi-FBS). Following mitogen withdrawal, cells were washed with 1X DPBS and incubated in media without mitogens and with low serum (RPMI, pen/strep, and 0.1% hi-FBS) for either 48 (2-day) or 96 (4-day) hours before harvesting. Contact inhibition was achieved by plating cells at 10% confluency (Lo) or 100% confluency (Hi) for 24 hours prior to switching to media with low serum (RPMI, pen/strep, and 0.1% hi-FBS) and with or without mitogens for 72 hours.

### Protein quantification and western blot analysis

For total protein quantification, 1 × 10^6^ FACS-isolated melb-a cells were lysed in RIPA buffer and total protein was quantified using a standard BCA protocol and Epoch spectrophotometer.

For western blot analysis, 1 × 10^6^ FACS-isolated melb-A cells were lysed in freshly prepared 1X Laemmli buffer at a concentration of 10,000 cells per µl. Samples were briefly sonicated and boiled at 95 °C for 5 minutes. Equal volumes of the sample were loaded onto a 12% polyacrylamide gel and electrophoresed at 100 volts for 1 hour before being transferred to a PVDF membrane using the iBlot 2 system (Thermo Fisher Scientific IB21001). Membrane blocking was performed at room temperature in 3% nonfat dried milk in PBST for 1 hour before incubating overnight with primary antibodies (1:1000 in PBST) at 4°C. After washing in PBST, the membrane was incubated with HRP-conjugated secondary antibodies (1:10,000 in PBST) for 1 hour at room temperature. Pierce ECL Western Blotting Substrate (Thermo Fisher Scientific 32106) was applied directly to the membrane for band detection and imaged using Bio-Rad ChemiDoc XRS+ System (Bio-Rad Laboratories, RRID:SCR_008426). Image analysis and densitometry of protein bands were performed using Image Lab software (v6.0.1, Image Lab Software, RRID:SCR_014210). Primary antibodies used for WB analysis included: anti-PDL1 (Abcam, ab269684, RRID:AB_2832197), anti-PCNA (Abcam, ab29, RRID:AB_303394), and anti-alpha-Smooth Muscle Actin (Thermo Fisher Scientific, PA5-16697, RRID:AB_11000908). Secondary antibodies employed were HRP conjugated, anti-mouse IgG (Thermo Fisher Scientific, 62-6520, RRID: AB_2533947), and anti-rabbit IgG (Cell Signaling Technology, 7074, RRID: AB_2099233).

### EdU assay

DNA synthesis was visualized using the flow cytometry EdU Click-iT assay (Thermo Fisher Scientific c10632) following manufacturer guidelines and carried out at room temperature and protected from light. Briefly, cells were incubated in media supplemented with 10 µM EdU for 2 hours at 37°C and 10% CO2 and washed in 1% BSA in 1X DPBS prior to harvesting. Cells were resuspended in 100 µL of Click-iT fixative for 15 minutes. Cells were washed and resuspended in 100 µL of 1X Click-iT saponin-based permeabilization and wash reagent for an additional 15 minutes before adding 0.5 mL of freshly prepared Click-iT reaction cocktail and incubated for 30 minutes. Cells were washed and resuspended in 500uL 1X Click-iT saponin-based permeabilization and wash reagent and immediately analyzed using an S3e BioRad cell sorter.

## Supplemental Information

Supplemental File 1: RNA-seq differential expression results comparing melanoblast (P0.5) to adult (8-week) McSCs and adult (8-week) McSCs to aged McSCs (24-month), along with core network information.

Supplemental File 2: Complete list of Hub Genes and TFs identified as having differential expression between melanoblast (P0.5) and adult (8-week) mice.

Supplemental File 3: Cross-sectional analysis of melanoblast (P0.5), adult (8-week) McSCs, and aged McSCs (24-month) DEGs including gene ontology analysis.

## References

Adelman, R.C., Stein, G., Roth, G.S., and Englander, D. (1972). Age-dependent regulation of mammalian DNA synthesis and cell proliferation In vivo. Mech. Ageing Dev. 1, 49–59. https://doi.org/10.1016/0047-6374(72)90052-8.

Agudo, J., Park, E.S., Rose, S.A., Alibo, E., Sweeney, R., Dhainaut, M., Kobayashi, K.S., Sachidanandam, R., Baccarini, A., Merad, M., et al. (2018). Quiescent Tissue Stem Cells Evade Immune Surveillance. Immunity 48, 271-285.e5. https://doi.org/10.1016/j.immuni.2018.02.001.

Ahmed, A.S.I., Sheng, M.H., Wasnik, S., Baylink, D.J., and Lau, K.-H.W. (2017). Effect of aging on stem cells. World J. Exp. Med. 7, 1–10. https://doi.org/10.5493/wjem.v7.i1.1.

Akinleye, A., and Rasool, Z. (2019). Immune checkpoint inhibitors of PD-L1 as cancer therapeutics. J. Hematol. Oncol.J Hematol Oncol 12, 92. https://doi.org/10.1186/s13045-019-0779-5.

Anderson, Z.T., Palmer, J.W., Idris, M.I., Villavicencio, K.M., Le, G., Cowart, J., Weinstein, D.E., and Harris, M.L. Topical RT1640 treatment effectively reverses gray hair and stem cell loss in a mouse model of radiation-induced canities. Pigment Cell Melanoma Res. n/a. https://doi.org/10.1111/pcmr.12913.

Augenlicht, L.H., and Baserga, R. (1974). Changes in the G0 state of WI-38 fibroblasts at different times after confluence. Exp. Cell Res. 89, 255–262. https://doi.org/10.1016/0014-4827(74)90789-7.

Bardhan, K., Anagnostou, T., and Boussiotis, V.A. (2016). The PD1:PD-L1/2 Pathway from Discovery to Clinical Implementation. Front. Immunol. 7. https://doi.org/10.3389/fimmu.2016.00550.

Basisty, N., Kale, A., Jeon, O.H., Kuehnemann, C., Payne, T., Rao, C., Holtz, A., Shah, S., Sharma, V., Ferrucci, L., et al. (2020). A proteomic atlas of senescence-associated secretomes for aging biomarker development. PLOS Biol. 18, e3000599. https://doi.org/10.1371/journal.pbio.3000599.

Baxter, L.L., Watkins-Chow, D.E., Pavan, W.J., and Loftus, S.K. (2018). A curated gene list for expanding the horizons of pigmentation biology. Pigment Cell Melanoma Res. https://doi.org/10.1111/pcmr.12743.

Bucher, N.L.R., and Swaffield, M.N. (1964). The Rate of Incorporation of Labeled Thymidine into the Deoxyribonucleic Acid of Regenerating Rat Liver in Relation to the Amount of Liver Excised. Cancer Res. 24, 1611–1625..

Bucher, N.L., Swaffield, M.N., and Ditroia, J.F. (1964). THE INFLUENCE OF AGE UPON THE INCORPORATION OF THYMIDINE-2-C14 INTO THE DNA OF REGENERATING RAT LIVER. Cancer Res. 24, 509–512..

Carter, L., Fouser, L.A., Jussif, J., Fitz, L., Deng, B., Wood, C.R., Collins, M., Honjo, T., Freeman, G.J., and Carreno, B.M. (2002). PD-1:PD-L inhibitory pathway affects both CD4(+) and CD8(+) T cells and is overcome by IL-2. Eur. J. Immunol. 32, 634–643. https://doi.org/10.1002/1521-4141(200203)32:3<634::AID-IMMU634>3.0.CO;2-9.

Casey, S.C., Tong, L., Li, Y., Do, R., Walz, S., Fitzgerald, K.N., Gouw, A.M., Baylot, V., Gütgemann, I., Eilers, M., et al. (2016). MYC regulates the antitumor immune response through CD47 and PD-L1. Science 352, 227–231. https://doi.org/10.1126/science.aac9935.

Chatterjee, A., Rodger, E.J., Ahn, A., Stockwell, P.A., Parry, M., Motwani, J., Gallagher, S.J., Shklovskaya, E., Tiffen, J., Eccles, M.R., et al. (2018). Marked Global DNA Hypomethylation Is Associated with Constitutive PD-L1 Expression in Melanoma. IScience 4, 312–325. https://doi.org/10.1016/j.isci.2018.05.021.

Chen, C.-C., Murray, P.J., Jiang, T.X., Plikus, M.V., Chang, Y.-T., Lee, O.K., Widelitz, R.B., and Chuong, C.-M. (2014). Regenerative Hair Waves in Aging Mice and Extra-Follicular Modulators Follistatin, Dkk1, and Sfrp4. J. Invest. Dermatol. 134, 2086–2096. https://doi.org/10.1038/jid.2014.139.

Chen, E.Y., Tan, C.M., Kou, Y., Duan, Q., Wang, Z., Meirelles, G.V., Clark, N.R., and Ma’ayan, A. (2013a). Enrichr: interactive and collaborative HTML5 gene list enrichment analysis tool. BMC Bioinformatics 14, 128. https://doi.org/10.1186/1471-2105-14-128.

Chen, S., Lewallen, M., and Xie, T. (2013b). Adhesion in the stem cell niche: biological roles and regulation. Dev. Camb. Engl. 140, 255–265. https://doi.org/10.1242/dev.083139.

Cheung, T.H., and Rando, T.A. (2013). Molecular regulation of stem cell quiescence. Nat. Rev. Mol. Cell Biol. 14. https://doi.org/10.1038/nrm3591.

Cheung, H.-H., Pei, D., and Chan, W.-Y. (2015). Stem cell aging in adult progeria. Cell Regen. 4. https://doi.org/10.1186/s13619-015-0021-z.

Chin, Y.E., Kitagawa, M., Su, W.-C.S., You, Z.-H., Iwamoto, Y., and Fu, X.-Y. (1996). Cell Growth Arrest and Induction of Cyclin-Dependent Kinase Inhibitor p21WAF1/CIP1 Mediated by STAT1. Science 272, 719–722. https://doi.org/10.1126/science.272.5262.719.

Cho, A.-R., Kim, J.Y., Munkhbayer, S., Shin, C.-Y., and Kwon, O. (2016). p21 upregulation in hair follicle stem cells is associated with telogen retention in aged mice. Exp. Dermatol. 25, 76–78. https://doi.org/10.1111/exd.12862.

Cho, I.J., Lui, P.P., Obajdin, J., Riccio, F., Stroukov, W., Willis, T.L., Spagnoli, F., and Watt, F.M. (2019). Mechanisms, Hallmarks, and Implications of Stem Cell Quiescence. Stem Cell Rep. 12, 1190–1200. https://doi.org/10.1016/j.stemcr.2019.05.012.

Coller, H.A., Sang, L., and Roberts, J.M. (2006). A New Description of Cellular Quiescence. PLOS Biol. 4, e83. https://doi.org/10.1371/journal.pbio.0040083.

Conboy, I.M., Conboy, M.J., Smythe, G.M., and Rando, T.A. (2003). Notch-Mediated Restoration of Regenerative Potential to Aged Muscle. Science 302, 1575–1577. https://doi.org/10.1126/science.1087573.

Coppé, J.-P., Desprez, P.-Y., Krtolica, A., and Campisi, J. (2010). The Senescence-Associated Secretory Phenotype: The Dark Side of Tumor Suppression. Annu. Rev. Pathol. 5, 99–118. https://doi.org/10.1146/annurev-pathol-121808-102144.

Dong, H., Zhu, G., Tamada, K., and Chen, L. (1999). B7-H1, a third member of the B7 family, co-stimulates T-cell proliferation and interleukin-10 secretion. Nat. Med. 5, 1365–1369. https://doi.org/10.1038/70932.

Du, J., Chen, Y., Li, Q., Han, X., Cheng, C., Wang, Z., Danielpour, D., Dunwoodie, S.L., Bunting, K.D., and Yang, Y.-C. (2012). HIF-1α deletion partially rescues defects of hematopoietic stem cell quiescence caused by Cited2 deficiency. Blood 119, 2789–2798. https://doi.org/10.1182/blood-2011-10-387902.

Fox, A.D., Hescott, B.J., Blumer, A.C., and Slonim, D.K. (2011). Connectedness of PPI network neighborhoods identifies regulatory hub proteins. Bioinformatics 27, 1135–1142. https://doi.org/10.1093/bioinformatics/btr099.

Freeman, G.J., Long, A.J., Iwai, Y., Bourque, K., Chernova, T., Nishimura, H., Fitz, L.J., Malenkovich, N., Okazaki, T., Byrne, M.C., et al. (2000). Engagement of the PD-1 immunoinhibitory receptor by a novel B7 family member leads to negative regulation of lymphocyte activation. J. Exp. Med. 192, 1027–1034. https://doi.org/10.1084/jem.192.7.1027.

Garcia-Diaz, A., Shin, D.S., Moreno, B.H., Saco, J., Escuin-Ordinas, H., Rodriguez, G.A., Zaretsky, J.M., Sun, L., Hugo, W., Wang, X., et al. (2017). Interferon Receptor Signaling Pathways Regulating PD-L1 and PD-L2 Expression. Cell Rep. 19, 1189–1201. https://doi.org/10.1016/j.celrep.2017.04.031.

Garrett, N.F.M. dos S., Costa, A.C.C. da, Ferreira, E.B., Damiani, G., Reis, P.E.D. dos, and Vasques, C.I. (2021). Prevalence of dermatological toxicities in patients with melanoma undergoing immunotherapy: Systematic review and meta-analysis. PLOS ONE 16, e0255716. https://doi.org/10.1371/journal.pone.0255716.

Gong, J., Chehrazi-Raffle, A., Reddi, S., and Salgia, R. (2018). Development of PD-1 and PD-L1 inhibitors as a form of cancer immunotherapy: a comprehensive review of registration trials and future considerations. J. Immunother. Cancer 6, 8. https://doi.org/10.1186/s40425-018-0316-z.

Goodell, M.A., and Rando, T.A. (2015). Stem cells and healthy aging. Science 350, 1199–1204. https://doi.org/10.1126/science.aab3388.

Gray, J.V., Petsko, G.A., Johnston, G.C., Ringe, D., Singer, R.A., and Werner-Washburne, M. (2004). “Sleeping Beauty”: Quiescence in Saccharomyces cerevisiae. Microbiol. Mol. Biol. Rev. 68, 187–206. https://doi.org/10.1128/MMBR.68.2.187-206.2004.

Guida, M., Strippoli, S., Maule, M., Quaglino, P., Ramondetta, A., Chiaron Sileni, V., Antonini Cappellini, G., Queirolo, P., Ridolfi, L., Del Vecchio, M., et al. (2021). Immune checkpoint inhibitor associated vitiligo and its impact on survival in patients with metastatic melanoma: an Italian Melanoma Intergroup study. ESMO Open 6, 100064. https://doi.org/10.1016/j.esmoop.2021.100064.

Harris, M.L., Fufa, T.D., Palmer, J.W., Joshi, S.S., Larson, D.M., Incao, A., Gildea, D.E., Trivedi, N.S., Lee, A.N., Day, C.-P., et al. (2018). A direct link between MITF, innate immunity, and hair graying. PLOS Biol. 16, e2003648. https://doi.org/10.1371/journal.pbio.2003648.

Hawkshaw, N.J., Hardman, J.A., Haslam, I.S., Shahmalak, A., Gilhar, A., Lim, X., and Paus, R. (2018). Identifying novel strategies for treating human hair loss disorders: Cyclosporine A suppresses the Wnt inhibitor, SFRP1, in the dermal papilla of human scalp hair follicles. PLOS Biol. 16, e2003705. https://doi.org/10.1371/journal.pbio.2003705.

Hirano, T., Honda, T., Kanameishi, S., Honda, Y., Egawa, G., Kitoh, A., Nakajima, S., Otsuka, A., Nomura, T., Dainichi, T., et al. (2021). PD-L1 on mast cells suppresses effector CD8+ T-cell activation in the skin in murine contact hypersensitivity. J. Allergy Clin. Immunol. 148, 563-573.e7. https://doi.org/10.1016/j.jaci.2020.12.654.

Horsley, V., Aliprantis, A.O., Polak, L., Glimcher, L.H., and Fuchs, E. (2008). NFATc1 Balances Quiescence and Proliferation of Skin Stem Cells. Cell 132, 299–310. https://doi.org/10.1016/j.cell.2007.11.047.

Hsu, J.-M., Li, C.-W., Lai, Y.-J., and Hung, M.-C. (2018). Posttranslational Modifications of PD-L1 and Their Applications in Cancer Therapy. Cancer Res. 78, 6349–6353. https://doi.org/10.1158/0008-5472.CAN-18-1892.

Iakova, P., Awad, S.S., and Timchenko, N.A. (2003). Aging Reduces Proliferative Capacities of Liver by Switching Pathways of C/EBPα Growth Arrest. Cell 113, 495–506. https://doi.org/10.1016/S0092-8674(03)00318-0.

Infarinato, N.R., Stewart, K.S., Yang, Y., Gomez, N.C., Pasolli, H.A., Hidalgo, L., Polak, L., Carroll, T.S., and Fuchs, E. (2020). BMP signaling: at the gate between activated melanocyte stem cells and differentiation. Genes Dev. 34, 1713–1734. https://doi.org/10.1101/gad.340281.120.

Inomata, K., Aoto, T., Binh, N.T., Okamoto, N., Tanimura, S., Wakayama, T., Iseki, S., Hara, E., Masunaga, T., Shimizu, H., et al. (2009). Genotoxic Stress Abrogates Renewal of Melanocyte Stem Cells by Triggering Their Differentiation. Cell 137, 1088–1099. https://doi.org/10.1016/j.cell.2009.03.037.

Iwai, Y., Ishida, M., Tanaka, Y., Okazaki, T., Honjo, T., and Minato, N. (2002). Involvement of PD-L1 on tumor cells in the escape from host immune system and tumor immunotherapy by PD-L1 blockade. Proc. Natl. Acad. Sci. 99, 12293–12297. https://doi.org/10.1073/pnas.192461099.

Janzen, V., Forkert, R., Fleming, H.E., Saito, Y., Waring, M.T., Dombkowski, D.M., Cheng, T., DePinho, R.A., Sharpless, N.E., and Scadden, D.T. (2006). Stem-cell ageing modified by the cyclin-dependent kinase inhibitor p16INK4a. Nature 443, 421–426. https://doi.org/10.1038/nature05159.

Johnson, S.C. (2018). Nutrient Sensing, Signaling and Ageing: The Role of IGF-1 and mTOR in Ageing and Age-Related Disease. Subcell. Biochem. 90, 49–97. https://doi.org/10.1007/978-981-13-2835-0_3.

Joshi, S.S., Tandukar, B., Pan, L., Huang, J.M., Livak, F., Smith, B.J., Hodges, T., Mahurkar, A.A., and Hornyak, T.J. (2019). CD34 defines melanocyte stem cell subpopulations with distinct regenerative properties. PLoS Genet. 15, e1008034. https://doi.org/10.1371/journal.pgen.1008034.

Kalamakis, G., Brüne, D., Ravichandran, S., Bolz, J., Fan, W., Ziebell, F., Stiehl, T., Catalá-Martinez, F., Kupke, J., Zhao, S., et al. (2019). Quiescence Modulates Stem Cell Maintenance and Regenerative Capacity in the Aging Brain. Cell 176, 1407-1419.e14. https://doi.org/10.1016/j.cell.2019.01.040.

Katsimpardi, L., Litterman, N.K., Schein, P.A., Miller, C.M., Loffredo, F.S., Wojtkiewicz, G.R., Chen, J.W., Lee, R.T., Wagers, A.J., and Rubin, L.L. (2014). Vascular and Neurogenic Rejuvenation of the Aging Mouse Brain by Young Systemic Factors. Science 344, 630–634. https://doi.org/10.1126/science.1251141.

Keir, M.E., Butte, M.J., Freeman, G.J., and Sharpe, A.H. (2008). PD-1 and Its Ligands in Tolerance and Immunity. Annu. Rev. Immunol. 26, 677–704. https://doi.org/10.1146/annurev.immunol.26.021607.090331.

Keyes, B.E., Segal, J.P., Heller, E., Lien, W.-H., Chang, C.-Y., Guo, X., Oristian, D.S., Zheng, D., and Fuchs, E. (2013). Nfatc1 orchestrates aging in hair follicle stem cells. Proc. Natl. Acad. Sci. 110, E4950–E4959. https://doi.org/10.1073/pnas.1320301110.

Krishnamurthy, J., Ramsey, M.R., Ligon, K.L., Torrice, C., Koh, A., Bonner-Weir, S., and Sharpless, N.E. (2006). p16 INK4a induces an age-dependent decline in islet regenerative potential. Nature 443, 453–457. https://doi.org/10.1038/nature05092.

Kuleshov, M.V., Jones, M.R., Rouillard, A.D., Fernandez, N.F., Duan, Q., Wang, Z., Koplev, S., Jenkins, S.L., Jagodnik, K.M., Lachmann, A., et al. (2016). Enrichr: a comprehensive gene set enrichment analysis web server 2016 update. Nucleic Acids Res. 44, W90–97. https://doi.org/10.1093/nar/gkw377.

Kwon, J.S., Everetts, N.J., Wang, X., Wang, W., Della Croce, K., Xing, J., and Yao, G. (2017). Controlling Depth of Cellular Quiescence by an Rb-E2F Network Switch. Cell Rep. 20, 3223–3235. https://doi.org/10.1016/j.celrep.2017.09.007.

Leeman, D.S., Hebestreit, K., Ruetz, T., Webb, A.E., McKay, A., Pollina, E.A., Dulken, B.W., Zhao, X., Yeo, R.W., Ho, T.T., et al. (2018). Lysosome activation clears aggregates and enhances quiescent neural stem cell activation during aging. Science 359, 1277–1283. https://doi.org/10.1126/science.aag3048.

Li, C.-W., Lim, S.-O., Xia, W., Lee, H.-H., Chan, L.-C., Kuo, C.-W., Khoo, K.-H., Chang, S.-S., Cha, J.-H., Kim, T., et al. (2016). Glycosylation and stabilization of programmed death ligand-1 suppresses T-cell activity. Nat. Commun. 7, 1–11. https://doi.org/10.1038/ncomms12632.

Liu, L., Cheung, T.H., Charville, G.W., Hurgo, B.M.C., Leavitt, T., Shih, J., Brunet, A., and Rando, T.A. (2013). Chromatin Modifications as Determinants of Muscle Stem Cell Quiescence and Chronological Aging. Cell Rep. 4, 189–204. https://doi.org/10.1016/j.celrep.2013.05.043.

Liu, Y., Gu, H.-Y., Zhu, J., Niu, Y.-M., Zhang, C., and Guo, G.-L. (2019). Identification of Hub Genes and Key Pathways Associated With Bipolar Disorder Based on Weighted Gene Co-expression Network Analysis. Front. Physiol. 10. https://doi.org/10.3389/fphys.2019.01081.

Lovén, J., Orlando, D.A., Sigova, A.A., Lin, C.Y., Rahl, P.B., Burge, C.B., Levens, D.L., Lee, T.I., and Young, R.A. (2012). Revisiting Global Gene Expression Analysis. Cell 151, 476–482. https://doi.org/10.1016/j.cell.2012.10.012.

Mak, S.-S., Moriyama, M., Nishioka, E., Osawa, M., and Nishikawa, S.-I. (2006). Indispensable role of Bcl2 in the development of the melanocyte stem cell. Dev. Biol. 291, 144–153. https://doi.org/10.1016/j.ydbio.2005.12.025.

Manson, G., Marabelle, A., and Houot, R. (2018). Hair Repigmentation With Anti–PD-1 and Anti–PD-L1 Immunotherapy: A Novel Hypothesis. JAMA Dermatol. 154, 113. https://doi.org/10.1001/jamadermatol.2017.4421.

Meng, Y., Liang, H., Hu, J., Liu, S., Hao, X., Wong, M.S.K., Li, X., and Hu, L. (2018). PD-L1 Expression Correlates With Tumor Infiltrating Lymphocytes And Response To Neoadjuvant Chemotherapy In Cervical Cancer. J. Cancer 9, 2938–2945. https://doi.org/10.7150/jca.22532.

Mourikis, P., Sambasivan, R., Castel, D., Rocheteau, P., Bizzarro, V., and Tajbakhsh, S. (2012). A critical requirement for notch signaling in maintenance of the quiescent skeletal muscle stem cell state. Stem Cells Dayt. Ohio 30, 243–252. https://doi.org/10.1002/stem.775.

Müller-Röver, S., Foitzik, K., Paus, R., Handjiski, B., van der Veen, C., Eichmüller, S., McKay, I.A., and Stenn, K.S. (2001). A Comprehensive Guide for the Accurate Classification of Murine Hair Follicles in Distinct Hair Cycle Stages. J. Invest. Dermatol. 117, 3–15. https://doi.org/10.1046/j.0022-202x.2001.01377.x.

Nishimura, E.K., Jordan, S.A., Oshima, H., Yoshida, H., Osawa, M., Moriyama, M., Jackson, I.J., Barrandon, Y., Miyachi, Y., and Nishikawa, S.-I. (2002). Dominant role of the niche in melanocyte stem-cell fate determination. Nature 416, 854–860. https://doi.org/10.1038/416854a.

Nishimura, E.K., Granter, S.R., and Fisher, D.E. (2005). Mechanisms of Hair Graying: Incomplete Melanocyte Stem Cell Maintenance in the Niche. Science 307, 720–724. https://doi.org/10.1126/science.1099593.

Nishimura, E.K., Suzuki, M., Igras, V., Du, J., Lonning, S., Miyachi, Y., Roes, J., Beermann, F., and Fisher, D.E. (2010). Key Roles for Transforming Growth Factor β in Melanocyte Stem Cell Maintenance. Cell Stem Cell 6, 130–140. https://doi.org/10.1016/j.stem.2009.12.010.

Oh, J., Lee, Y.D., and Wagers, A.J. (2014). Stem cell aging: mechanisms, regulators and therapeutic opportunities. Nat. Med. 20, 870–880. https://doi.org/10.1038/nm.3651.

Paus, R., Ito, N., Takigawa, M., and Ito, T. (2003). The Hair Follicle and Immune Privilege. J. Investig. Dermatol. Symp. Proc. 8, 188–194. https://doi.org/10.1046/j.1087-0024.2003.00807.x.

Peters, E.M.J., Maurer, M., Botchkarev, V.A., Jensen, K. deMasey, Welker, P., Scott, G.A., and Paus, R. (2003). Kit Is Expressed by Epithelial Cells In Vivo. J. Invest. Dermatol. 121, 976–984. https://doi.org/10.1046/j.1523-1747.2003.12478.x.

Qin, W., Hu, L., Zhang, X., Jiang, S., Li, J., Zhang, Z., and Wang, X. (2019). The Diverse Function of PD-1/PD-L Pathway Beyond Cancer. Front. Immunol. 10..

Rabbani, P., Takeo, M., Chou, W., Myung, P., Bosenberg, M., Chin, L., Taketo, M.M., and Ito, M. (2011). Coordinated Activation of Wnt in Epithelial and Melanocyte Stem Cells Initiates Pigmented Hair Regeneration. Cell 145, 941–955. https://doi.org/10.1016/j.cell.2011.05.004.

Raedler, L.A. (2015). Keytruda (Pembrolizumab): First PD-1 Inhibitor Approved for Previously Treated Unresectable or Metastatic Melanoma. Am. Health Drug Benefits 8, 96–100..

Rivera, N., Boada, A., Bielsa, M.I., Fernández-Figueras, M.T., Carcereny, E., Moran, M.T., and Ferrándiz, C. (2017). Hair Repigmentation During Immunotherapy Treatment With an Anti– Programmed Cell Death 1 and Anti–Programmed Cell Death Ligand 1 Agent for Lung Cancer. JAMA Dermatol. 153, 1162–1165. https://doi.org/10.1001/jamadermatol.2017.2106.

Rodgers, J.T., King, K.Y., Brett, J.O., Cromie, M.J., Charville, G.W., Maguire, K.K., Brunson, C., Mastey, N., Liu, L., Tsai, C.-R., et al. (2014). mTORC1 controls the adaptive transition of quiescent stem cells from G0 to GAlert. Nature 510, 393–396. https://doi.org/10.1038/nature13255.

Rumman, M., Dhawan, J., and Kassem, M. (2015). Concise Review: Quiescence in Adult Stem Cells: Biological Significance and Relevance to Tissue Regeneration: ASC Quiescence: Role and Relevance to Tissue Regeneration. STEM CELLS 33, 2903–2912. https://doi.org/10.1002/stem.2056.

Salic, A., and Mitchison, T.J. (2008). A chemical method for fast and sensitive detection of DNA synthesis in vivo. Proc. Natl. Acad. Sci. 105, 2415–2420. https://doi.org/10.1073/pnas.0712168105.

Sharifi, S., da Costa, H.F.R., and Bierhoff, H. (2020). The circuitry between ribosome biogenesis and translation in stem cell function and ageing. Mech. Ageing Dev. 189, 111282. https://doi.org/10.1016/j.mad.2020.111282.

Slominski, A., Paus, R., Plonka, P., Chakraborty, A., Maurer, M., Pruski, D., and Lukiewicz, S. (1994). Melanogenesis During the Anagen-Catagen-Telogen Transformation of the Murine Hair Cycle. J. Invest. Dermatol. 102, 862–869. https://doi.org/10.1111/1523-1747.ep12382606.

Slominski, A., Wortsman, J., Plonka, P.M., Schallreuter, K.U., Paus, R., and Tobin, D.J. (2005). Hair Follicle Pigmentation. J. Invest. Dermatol. 124, 13–21. https://doi.org/10.1111/j.0022-202X.2004.23528.x.

Sviderskaya, E.V., Wakeling, W.F., and Bennett, D.C. (1995). A cloned, immortal line of murine melanoblasts inducible to differentiate to melanocytes. Development 121, 1547–1557..

Szklarczyk, D., Gable, A.L., Lyon, D., Junge, A., Wyder, S., Huerta-Cepas, J., Simonovic, M., Doncheva, N.T., Morris, J.H., Bork, P., et al. (2019). STRING v11: protein–protein association networks with increased coverage, supporting functional discovery in genome-wide experimental datasets. Nucleic Acids Res. 47, D607–D613. https://doi.org/10.1093/nar/gky1131.

Tanimura, S., Tadokoro, Y., Inomata, K., Binh, N.T., Nishie, W., Yamazaki, S., Nakauchi, H., Tanaka, Y., McMillan, J.R., Sawamura, D., et al. (2011). Hair Follicle Stem Cells Provide a Functional Niche for Melanocyte Stem Cells. Cell Stem Cell 8, 177–187. https://doi.org/10.1016/j.stem.2010.11.029.

Tümpel, S., and Rudolph, K.L. (2019). Quiescence: Good and Bad of Stem Cell Aging. Trends Cell Biol. 29, 672–685. https://doi.org/10.1016/j.tcb.2019.05.002.

Ueno, M., Aoto, T., Mohri, Y., Yokozeki, H., and Nishimura, E.K. (2014). Coupling of the radiosensitivity of melanocyte stem cells to their dormancy during the hair cycle. Pigment Cell Melanoma Res. 27, 540–551. https://doi.org/10.1111/pcmr.12251.

Uno, M., and Nishida, E. (2016). Lifespan-regulating genes in C. elegans. Npj Aging Mech. Dis. 2, 1–8. https://doi.org/10.1038/npjamd.2016.10.

Urbán, N., and Cheung, T.H. (2021). Stem cell quiescence: the challenging path to activation. Dev. Camb. Engl. 148, dev165084. https://doi.org/10.1242/dev.165084.

van Veelen, W., Korsse, S.E., van de Laar, L., and Peppelenbosch, M.P. (2011). The long and winding road to rational treatment of cancer associated with LKB1/AMPK/TSC/mTORC1 signaling. Oncogene 30, 2289–2303. https://doi.org/10.1038/onc.2010.630.

Velthoven, C.T.J. van, and Rando, T.A. (2019). Stem Cell Quiescence: Dynamism, Restraint, and Cellular Idling. Cell Stem Cell 24, 213–225. https://doi.org/10.1016/j.stem.2019.01.001.

Yale, K., Juhasz, M., and Atanaskova Mesinkovska, N. (2020). Medication-Induced Repigmentation of Gray Hair: A Systematic Review. Skin Appendage Disord. 6, 1–10. https://doi.org/10.1159/000504414.

Yao, Y., Tao, R., Wang, X., Wang, Y., Mao, Y., and Zhou, L.F. (2009). B7-H1 is correlated with malignancy-grade gliomas but is not expressed exclusively on tumor stem-like cells. Neuro-Oncol. 11, 757–766. https://doi.org/10.1215/15228517-2009-014.

Yi, M., Niu, M., Xu, L., Luo, S., and Wu, K. (2021). Regulation of PD-L1 expression in the tumor microenvironment. J. Hematol. Oncol. J Hematol Oncol 14, 10. https://doi.org/10.1186/s13045-020-01027-5.

Zhang, J., Bu, X., Wang, H., Zhu, Y., Geng, Y., Nihira, N.T., Tan, Y., Ci, Y., Wu, F., Dai, X., et al. (2018). Cyclin D–CDK4 kinase destabilizes PD-L1 via cullin 3–SPOP to control cancer immune surveillance. Nature 553, 91–95. https://doi.org/10.1038/nature25015.

Zhang, L., Gajewski, T.F., and Kline, J. (2009). PD-1/PD-L1 interactions inhibit antitumor immune responses in a murine acute myeloid leukemia model. Blood 114, 1545–1552. https://doi.org/10.1182/blood-2009-03-206672.

Zhang, N., Zeng, Y., Du, W., Zhu, J., Shen, D., Liu, Z., and Huang, J.-A. (2016). The EGFR pathway is involved in the regulation of PD-L1 expression via the IL-6/JAK/STAT3 signaling pathway in EGFR-mutated non-small cell lung cancer. Int. J. Oncol. 49, 1360–1368. https://doi.org/10.3892/ijo.2016.3632.

Zhou, L., Wen, L., Sheng, Y., Lu, J., Hu, R., Wang, X., Lu, Z., and Yang, Q. (2021). The PD-1/PD-L1 pathway in murine hair cycle transition: a potential anagen phase regulator. Arch. Dermatol. Res. 313, 751–758. https://doi.org/10.1007/s00403-020-02169-9.

Zhou, Y., Yan, X., Feng, X., Bu, J., Dong, Y., Lin, P., Hayashi, Y., Huang, R., Olsson, A., Andreassen, P.R., et al. (2018). Setd2 regulates quiescence and differentiation of adult hematopoietic stem cells by restricting RNA polymerase II elongation. Haematologica 103, 1110–1123. https://doi.org/10.3324/haematol.2018.187708.

